# Caste- and Sex-Specific Differential Investment in Brain Regions of Australian Ants

**DOI:** 10.1101/2025.09.23.678185

**Authors:** Saroja Ellendula, Zachary BV Sheehan, Marcel E Sayre, Fleur Ponton, Ajay Narendra

## Abstract

Even within a single species, closely related individuals exhibit distinct lifestyles that demand different information processing requirements. Worker ants are ambulatory and handle tasks like colony maintenance and foraging, while winged reproductive castes focus on mating and colony founding. Here, we compare the volumes of functionally distinct brain regions across alate males, alate females, and workers in two species of ants native to Australia, *Myrmecia midas* and *Rhytidoponera metallica*, to assess adaptations to behavioural, ecological, and locomotor demands. Female castes in both species had larger brains with pronounced mushroom bodies, supporting their broader behavioural repertoire and navigational tasks. In comparison, males exhibited smaller brains but enlarged optic lobes and central complexes, highlighting the significance of vision and orientation in mate searching. Species-specific patterns were also noted: *R. metallica* individuals consistently had larger antennal lobes and smaller optic lobes across castes, indicating a reliance on olfactory cues. However, the mushroom bodies remained comparatively similar. These findings demonstrate how caste- and species-specific sensory demands shape neural architecture and serve as a basis for understanding the interplay between brain structure and function.

## Introduction

Insects offer an unparalleled opportunity to explore the neural basis of behaviour and cognition in an evolutionary context. Though an insect brain weighs a milligram and contains orders of magnitude fewer neurons than a vertebrate brain, these miniaturised brains support sophisticated navigation, communication, decision-making, spatial memory and learning. Compartmentalised organisation of insect brains into discrete neuropils, which are functionally specialised processing centres, provides an ideal system to investigate how evolutionary pressure shapes neural investment strategy (Bouchebti and Arganda, 2020; Rozanski et al., 2022).

Insect brains exhibit remarkable plasticity throughout an individual’s lifetime, with neuropil volumes, cell numbers and connectivity changing in response to modifications in lifestyle and information content in the environment. For instance, butterflies that show complex foraging behaviours and enhanced long-term memory have larger neuropils involved in learning and memory (mushroom bodies) when compared to butterflies that exhibit simpler behavioural repertoires (Couto et al., 2023; Farnworth et al., 2024). Similarly, experienced foragers tend to have larger mushroom bodies compared to similarly aged animals that stay within the confines of the nest (Gronenberg et al., 1996; Kühn-Bühlmann and Wehner, 2006; Riveros and Gronenberg, 2010; Withers et al., 1993). As insects age and are exposed to light, there is a significant reduction of the synaptic complexes (microglomeruli) in this region, whereas associative learning and long-term memory formation promote local increases in microglomerular number; in both cases, these synaptic changes are accompanied by increased dendritic branching of intrinsic mushroom body neurons (kenyon cells; Cabirol et al., 2018; Farris et al., 2001; Grob et al., 2019; Grob et al., 2017; Groh and Rössler, 2020; Narasimhan et al., 2025; Scholl et al. 2014; Steib et al., 2010; Seid and Wehner 2009; Yilmaz et al., 2016). Beyond age and experience, shifts in lifestyle can also reshape insect brain architecture. In locusts, for example, the central complex, a neuropil associated with goal-directed movement (Honkanen et al., 2019), significantly increases in volume when animals transition from a solitary to a gregarious phase, an adaptation likely to assist movement of individual animals within a dense swarm and to deal with intense competition (Ott and Rogers, 2010). Such plasticity underscores the capacity of an insect brain to match form to function across different lifestyles and ecological contexts.

Ants provide a unique opportunity to study the link between brain, behaviour and ecology within a species, due to the distinct lifestyles exhibited by the different castes. Workers, the most numerous caste, are typically sterile females who are exclusively ambulatory and perform the largest share of tasks within the colony, including brood care, nest maintenance and foraging. For successful foraging, workers rely on different sensory modalities to locate food sources and to find their way back home (Collett et al., 2025). In contrast, the reproductive castes, alate males and females, have a limited behavioural repertoire focused primarily on mating and colony founding (Gronenberg, 2008). Alate males and females rely on visual information for flight control, navigation, and obstacle avoidance. Males also use visual cues to track fast-moving conspecific females and fend off competitors. While males die after mating, alate females shed their wings and establish new nests as queens, spending the rest of their lives in the darkness. In some species (e.g., *Myrmecia*), queens forage and return to their newly found colony until the first brood of workers emerges (Reid et al., 2013; Haskins and Haskins, 1950).

Variation in lifestyle between castes and species is reflected in the organisation of their external sensory array. Male ants tend to have disproportionately large compound eyes that contain smaller and more lenses in each eye compared to any other caste (Gronenberg and Hölldobler, 1999; Narendra et al., 2011; Narendra et al., 2016). Among all castes, males possess the largest simple eyes, known as ocelli, which aid in altitude control, horizon detection, and in detecting changes in the polarised skylight pattern (Narendra and Ribi, 2017; Penmetcha et al., 2019). Across all ants, nocturnal species typically have larger lenses and wider photoreceptors to capture more light (Greiner et al., 2007; Moser et al., 2004; Narendra et al., 2011; Narendra et al., 2017). Caste- and sex-specific differences were also found in external antennal morphology, critical for chemical communication. In *Camponotus japonicus*, alate females had the most sensilla and males had the least (Nakanishi et al., 2009). While certain types of sensilla were common across all castes, their distribution on the antennae differed significantly, with one type of sensilla (basiconica sensillum) noticeably absent in male antennae (Nakanishi et al., 2009).

Information captured from the external sensory structures is transmitted via sensory neurons to the primary sensory neuropils (optic lobes, antennal lobes) and subsequently to higher-order information-processing centres (mushroom bodies, central complex). The volume of the neuropils provides a glimpse into the relative investment an animal has made into processing a specific type of sensory information, supporting associative memory, or mediating sensorimotor transformations. Nocturnal hawkmoths and paper wasps, for example, invested more in their antennal lobes compared to optic lobes, suggesting less reliance on visual information (Stöckl et al., 2016; O’Donnell et al., 2013). Cave beetles that spend their entire lives in complete darkness have evolved to lose their visual processing centres altogether (Ghaffar et al., 1984). Among worker ants of *Myrmecia* active at discrete times of the day, nocturnal ants invested less in primary visual processing neuropils and more in primary olfactory processing regions and downstream neuropils that integrate visual and olfactory information (Sheehan et al., 2019).

In this study, we aimed to identify the differential investment in brain regions between distinct castes and sexes in two visually oriented solitary foraging Australian ants, *Myrmecia midas* and *Rhytidoponera metallica*. Both species exhibited dramatic variation in body size and in their activity times. Workers of *M. midas* are strictly nocturnal, but the winged males and females fly out during the day, which leads to distinct activity times of castes within a single species (pers. obs. A.N.; see also Narendra et al., 2011). In contrast, in *R. metallica*, all three castes are strictly day-active. Workers of *M. midas* were more than twice the size of *R. metallica*. Workers of both species use visual cues for navigation and obtain compass information from celestial cues (Freas et al., 2024; Joseph, 2023). Young queens of both species (*R. metallica*: Ward, 1986; *M. midas*: Reid et al., 2013; Haskins and Haskins, 1950) also spend a brief period in their life foraging while establishing new nests. We analysed the brains of workers, alate females and males in both ant species and carried out a scaling analysis of distinct neuropils.

## Methods

### Study species and location

We studied workers, alate males and alate females of *Rhytidoponera metallica* (n = 7, 6, and 8, respectively) and alate males and alate females of *Myrmecia midas* (n = 7 and 6, respectively). Data for *M. midas* workers (n = 28) were obtained from Sheehan et al. (2019; details in the section ‘Data for *M. midas* workers’). Ants were collected from multiple nests at the Macquarie University Wallumattagal Campus in Sydney, Australia (33°46’10.24” S, 151°06’39.55” E) between 2017 and 2023. Workers were collected at the nest entrance or while foraging at different times throughout the year. Alate females and males were collected between late January and mid-April as they left the nest entrance for nuptial flights. Captured animals were immediately dissected, and the samples were processed.

### Immunohistochemistry and imaging

We processed the samples using the same protocol as described by Sheehan et al. (2019). Animals were anaesthetised on ice before removing the brain from the head capsule. The dorsal surface of the head was photographed with a colour camera (Lumix DMC-FZ1000, Panasonic Australia). From these images, the individual’s head width (HW) was determined by measuring along the widest point of the head.

Brains were dissected from the head capsule in Ringer solution (129 mM NaCl, 6 mM KCl, 4.3 mM MgCl_2_ × 6H_2_O, 5 mM CaCl_2_ × 2H_2_O, 159.8 mM Sucrose, 274 mM D-glucose, 10 mM HEPES buffer, pH 6.7-7). Brains were transferred within 10 minutes to a fixative: 4% paraformaldehyde (PFA) in 0.1 M Phosphate Buffered Solution (PBS). Brains were kept on a shaker in the dark in all the following described washes and incubations. Brains were kept in the fixative for two days at room temperature and washed three times (10 minutes each) in PBS. To assist antibody penetration, specimens were washed with 3% Triton-X in 0.1M PBS (PBST) at room temperature. They were then incubated for one hour with 2% Normal Goat Serum (NGS, Sigma-Aldrich) in PBST. Brains were then incubated in the primary antibody solution, 1:50 anti-synapsin (3C11 anti SYNORF1; DHSB) for four days in 2% NGS in PBST, at room temperature. This was followed by five washes in PBS (10 minutes each). They were then incubated in the secondary antibody solution, 1:250 goat anti-mouse antibody (Alexa Flour 488 goat anti-mouse, ThermoFisher Scientific), for three days in 1% NGS in PBST, at room temperature. Following this, the brains were washed five times with 0.1 M PBS (10 minutes each) and dehydrated through an ascending series of ethanol (30%, 50%, 70%, 90%, 95% and 100%; 10 minutes each). Brains were cleared for 10 minutes in 1:1 methyl salicylate:ethanol, followed by 100% methyl salicylate (Sigma Aldrich, M6752) for one hour at room temperature. Lastly, brains were transferred to custom-made metal slides containing 1 cm diameter holes. A well was created by sealing the holes with a coverslip on one side. Brains were immersed in 100% methyl salicylate with the ventral side facing upward, and the well was sealed with another coverslip.

Two of the *M. midas* alate female brain samples were collected in 2023 to increase the sample size. These two brains were processed using the above-described protocol, with slight modifications. Following dissection, brains were fixed overnight at 4°C. Additionally, they were incubated in the primary and secondary antibody (1% NGS in PBST) for only two days each. These changes to the original protocol did not appear to have an impact on the fixation or the staining of the tissue. Lastly, to preserve specimens, these brains were mounted in 100% Permount.

All specimens (except the brains of the two *M. midas* alate females processed in 2023) were imaged using an inverted confocal laser scanning microscope (Olympus FluoView FV 1000 IX81) using a 10x objective (UPlanApo, NA 0.4) and 1-3.1 μm optical sections. The two specimens processed in 2023 were imaged using an Olympus FluoView FV 3000RS IX83 confocal microscope with a 10x objective (UPLSAPO10X2). For ants with large brain size (seen in *M. midas*), we imaged three overlapping z-stacks that were merged together using the “Pairwise Stitching” plugin in the Fiji program (Schindelin et al. 2012; RRID:SCR_02285) to produce a single image.

### Data analyses

#### Brain volumes

Three-dimensional reconstructions of neuropils were done based on the anti-synapsin labelling to obtain volumes of individual regions. Neuropil boundaries were delineated in every third image in each image stack using Amira (v. 6.0.1, ThermoFisher Scientific). Two methods were used to produce complete representations of these regions: the interpolate function in Amira and a semi-automated interpolation method using ‘Biomedisa’ (Lösel et al. 2020). These interpolated contours were double-checked and manually corrected in cases where deviations from the neuropil boundaries were noted. The spherical aberration that is known to occur due to refractive index (RI) mismatch between air (RI = 1) and the mounting media was corrected by multiplying the optical section thickness by 1.55 and 1.54 when samples were mounted in methyl salicylate (RI = 1.536) and permount (RI = 1.520), respectively (calculated from Diel et al. 2020). The volume of the outlined regions was calculated using the “MaterialStatistics” function in Amira. Finally, 3D reconstructions were made using a polygonal surface model in Amira.

Four functionally defined areas with clear glial borders were quantified in each brain. These included the optic lobes (OL; containing the Lamina (LA), Medulla (ME) and Lobula (LO)), the Antennal lobe (AL), the Mushroom Body (MB; containing the Calyx Lip (LIP), Calyx Collar (COL) and the Peduncle (PED)) and the Central Complex (CX; containing the fan-shaped body (FB; also referred to as the upper division of the Central Body CBU), the ellipsoid body (EB; also referred to as the lower division of the Central Body CBL), the protocerebral bridge (PB) and the paired Noduli (NO); Figure 1). In our image data, the basal ring could not be reliably distinguished from the COL. Thus, ‘COL’ in this paper refers to the combined volume of the COL and the basal ring. The rest of the brain is composed of undefined neuropil mass and was thus segmented as one single region: the rest of the central brain (RoCB; which includes the unstructured lateral protocerebrum and the subesophageal ganglion). Paired neuropils were traced in only one hemisphere. This volume was multiplied by two to estimate the total volume of bilateral neuropils. The total neuropil volume of the brain was obtained by summing the volume of the paired and unpaired neuropils. No cell bodies were included in the measurements.

**Figure 1:**
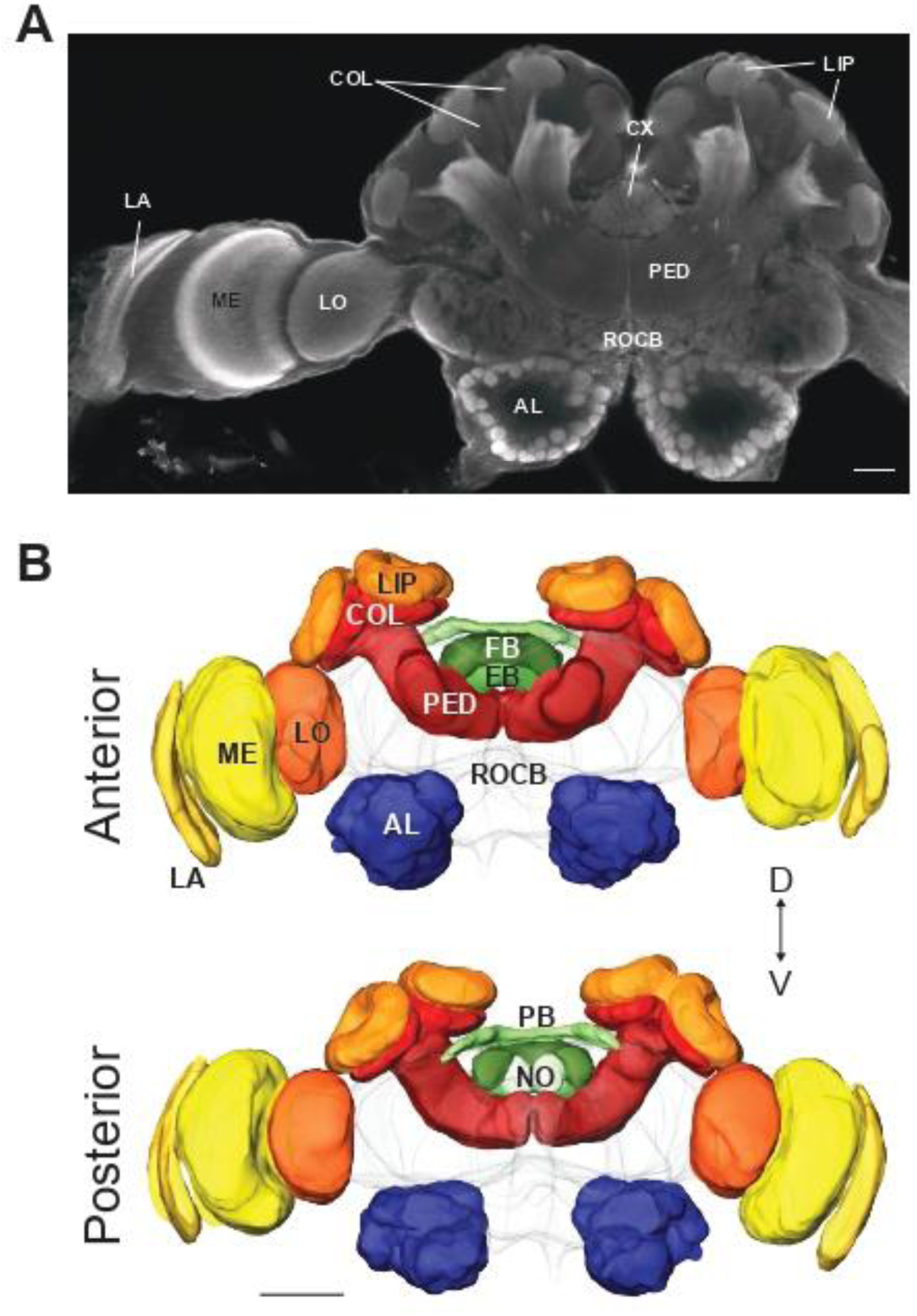
General layout of the brain in *Myrmecia midas* and *Rhytidoponera metallica*. (A) Frontal section of a *M. midas* worker brain labelled with anti-synapsin highlighting the major neuropils included in this study: the optic lobes (OL: lamina [LA], medulla [ME], lobula [LO]), antennal lobes (AL), mushroom bodies (MB: lip [LIP], collar [COL], peduncle [PED]), central complex (CX: fan-shaped body [FB], ellipsoid body [EB], protocerebral bridge [PB], noduli [NO]), and the rest of the central brain (RoCB) (B) Anterior (top panel) and posterior (bottom panel) views of surface reconstruction of neuropils in a representative male *R. metallica* brain. Scale bar = 100 μm.

#### Data for *M. midas* workers

For workers of *M. midas*, we used the raw image data and label files of the brains (n = 28) from Sheehan et al. (2019). Neuropil volumes were calculated as described earlier to ensure comparability with data collected as part of this study. Thus, the spherical aberration of the obtained data was corrected by multiplying the raw optical section thickness by 1.55, as samples in Sheehan et al. (2019) were also mounted in an identical methyl salicylate medium (Sigma Aldrich, M6752; RI = 1.536).

#### Statistical analyses

The two axes of comparison in this study were the castes within the species and the two species themselves. To investigate whether differences in neuropil volume between any groups were attributable to genuine differences in neuropil sizes among distinct groups rather than variations caused by allometric scaling, we performed a standardised major axis regression (SMA) analysis on individual neuropils in relation to the unstructured brain volume, the RoCB, using the SMATR (Warton et al., 2012) package in R (version 4.3.0). This analysis assumes an allometric relationship described as y = α · x to the power b, which can be translated to the linear relationship log(y) = log(x)·b+log(α). If the allometric scaling or the slopes between the tested groups were not statistically different, we tested for differences in the y-axis intercept or elevation, which implies a true difference in neuropil volumes across groups. Thus, all comparisons between groups using SMA refer to differences in neuropil volume relative to the reference neuropil volume, i.e., the RoCB. Lastly, a grade shift index (gsi), described as α1/α2 (Ott and Rogers, 2010), was calculated to estimate the magnitude of the shift in elevation, where ‘1’ and ‘2’ represent the two groups being compared.

## Results

### Variation in total neuropil volumes

Both the head width and total neuropil volume varied between all three castes of *M. midas* and *R. metallica* (Figure 2). Across all three castes, *M. midas* had a greater head width compared to *R. metallica*. Within a single species, males consistently had the smallest head size, and alate females were the largest. These trends were reflected in the head widths of the individuals (means ± s.d.; *M. midas*: workers = 3.45 ± 0.5 mm, alate females = 4.60 ± 0.07 mm, alate males = 2.61 ± 0.09 mm; *R. metallica*: workers = 1.53 ± 0.12 mm, alate females = 1.63 ± 0.07 mm, alate males = 1.24 ± 0.06 mm; Supplementary Table 1).

**Figure 2:**
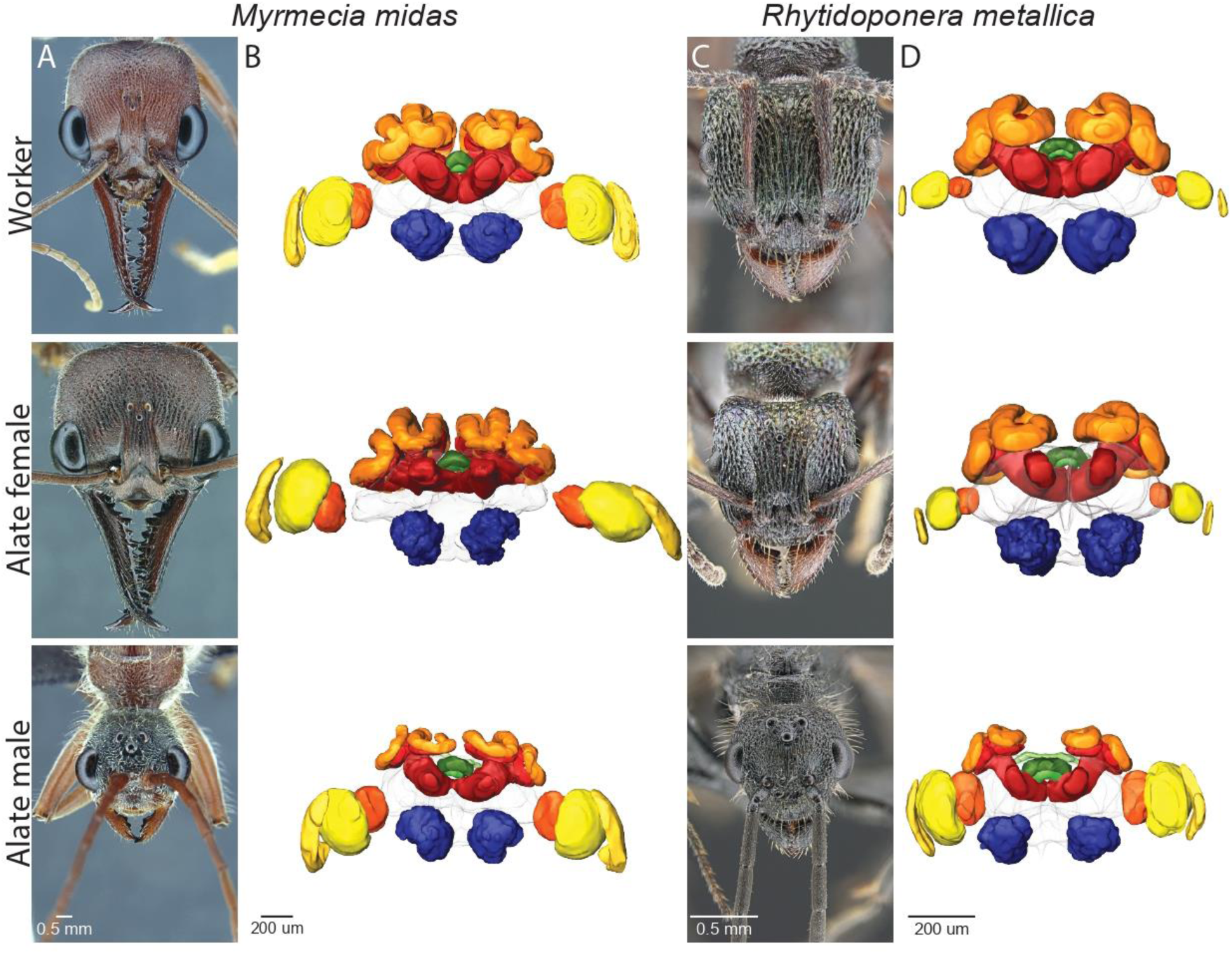
Comparative head and brain morphology of three castes in *Myrmecia midas* and *Rytidoponera metallica*. (A) and (C) Example images of the heads of a worker (top), an alate female (middle) and a male (bottom) ant to show variation in size and head morphology between castes in *M. midas* and *R. metallica*, respectively. (B) and (D) Representative surface renderings of a worker (top), an alate female (middle) and a male (bottom) brain in *M. midas* and *R. metallica*, respectively.

A similar trend was seen in the total absolute neuropil volume. Across all three castes, individuals of *M. midas* had larger total neuropil volume compared to *R. metallica* (by an order of magnitude), which correlated with their head widths. In both species, males had the smallest total neuropil volume and alate females the largest (mean±s.d.; *M. midas*: workers = 3.41 × 10^8^ ± 5.07 × 10^7^ µm^3^, alate females = 3.67 × 10^8^ ± 1.53 × 10^8^ μm^3^, alate males = 2.49 × 10^8^ ± 2.60 × 10^7^ μm^3^; *R. metallica*: workers = 3.97 × 10^7^ ± 1.09 × 10^7^ μm^3^, alate females = 4.25 × 10^7^ ± 3.88 × 10^6^ μm^3^, alate males = 2.37 × 10^7^ ± 1.16 × 10^7^ μm^3^; Supplementary Table 1).

### Optic Lobes (OL)

The OLs, comprising the LA, ME, and LO, are the primary visual processing centres in the insect brain (Figure 1). We first compared the scaling pattern of OL of each caste between species.

#### Comparison between species

In worker ants that are strictly ambulatory, the OL volume scaled differently in both *M. midas* and *R. metallica*, making it theoretically invalid to test for differences in a grade shift (Figure 3A, top panel). However, each of the three sub-neuropils of the OL scaled similarly, i.e., their slopes were not significantly different, and thus could be compared for differences in volume. The LA did not differ significantly between workers of the two species. However, relative to the reference structure, the RoCB, both the ME and LO, were significantly larger in the workers of *M. midas* compared to *R. metallica* (Figure 3A, top panel inset).

**Figure 3:**
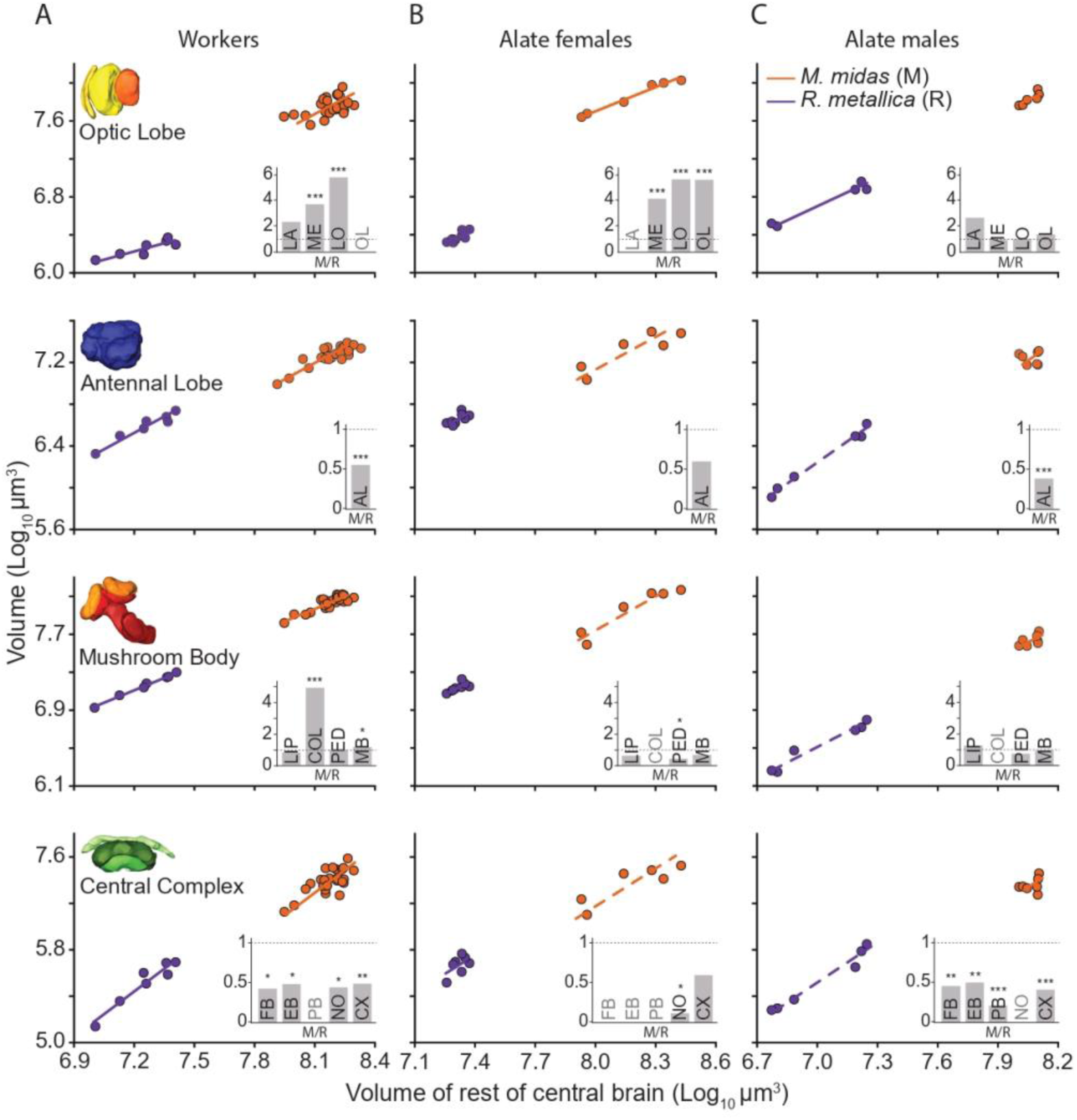
Scaling relationship of the volume of the major neuropils and the sub-neuropils between the two species, *Myrmecia midas* and *Rytidoponera metallica.* Comparison of scaling relations of the main neuropils to the reference structure, RoCB, between the two study species, *M. midas* (orange) and *R. metallica* (purple), across workers, alate females and alate males are shown. Each panel shows comparisons between species for the same neuropil. Regression lines are derived from SMA analysis and fitted to the data. Dashed lines represent non-significant linear regression models. The inset shows gsi for the region of interest when comparing both species. The gsi estimates the difference in neuropil size of the same caste between two species, calculated using the elevation of the regression line. From left to right, bars represent gsi for each sub-neuropil and the whole neuropil when comparing *M. midas* (M) to *R. metallica* (R). A gsi of 1 indicates that both species have a similarly sized neuropil, shown by the horizontal dotted line. When comparing both species, a gsi <1 indicates that the neuropil is larger in *R. metallica* than in *M. midas* in the comparison and vice versa. In cases where it was theoretically invalid to test for volumetric differences, no gsi was calculated (indicated by grey text).

In the alate females that lead a walking and flying lifestyle, relative to the RoCB, the OLs were larger in *M. midas* compared to *R. metallica* (Figure 3B, top panel). A volumetric difference was also found in the ME and LO, while the LA exhibited distinct scaling patterns in both species (Figure 3B, top panel inset).

Males that lead a life exclusively on the wing exhibited no significant differences in the volume of the OL (Figure 3C, top panel) and all sub-neuropils relative to the RoCB between the two species (Figure 3C, top panel inset; Supplementary Table 2).

#### Comparison within species

We compared the scaling relationships of neuropil volumes of the three castes within each species. In both *M. midas* and *R. metallica*, males exhibited significantly larger OL volumes relative to the RoCB than either of the female castes. In both species, between the two female castes, the alates possessed relatively larger OLs compared to the workers (Figure 4A-B, top panel). The magnitude of this difference between males and either of the female castes was more pronounced in *R. metallica* than in *M. midas* (Figure 4B, top panel inset). Male ants thus have the largest OL neuropil volume, while workers have the smallest, relative to RoCB. This pattern was consistent across all sub-neuropils of the OL (Figure 4), with few variations. For instance, in *M. midas*, males and alate females had comparable LA volumes (Figure 4A, second panel inset), while in *R. metallica*, alate females and workers shared similar LA and ME volumes (Figure 4B, second and third panel insets; Supplementary Table 3).

**Figure 4:**
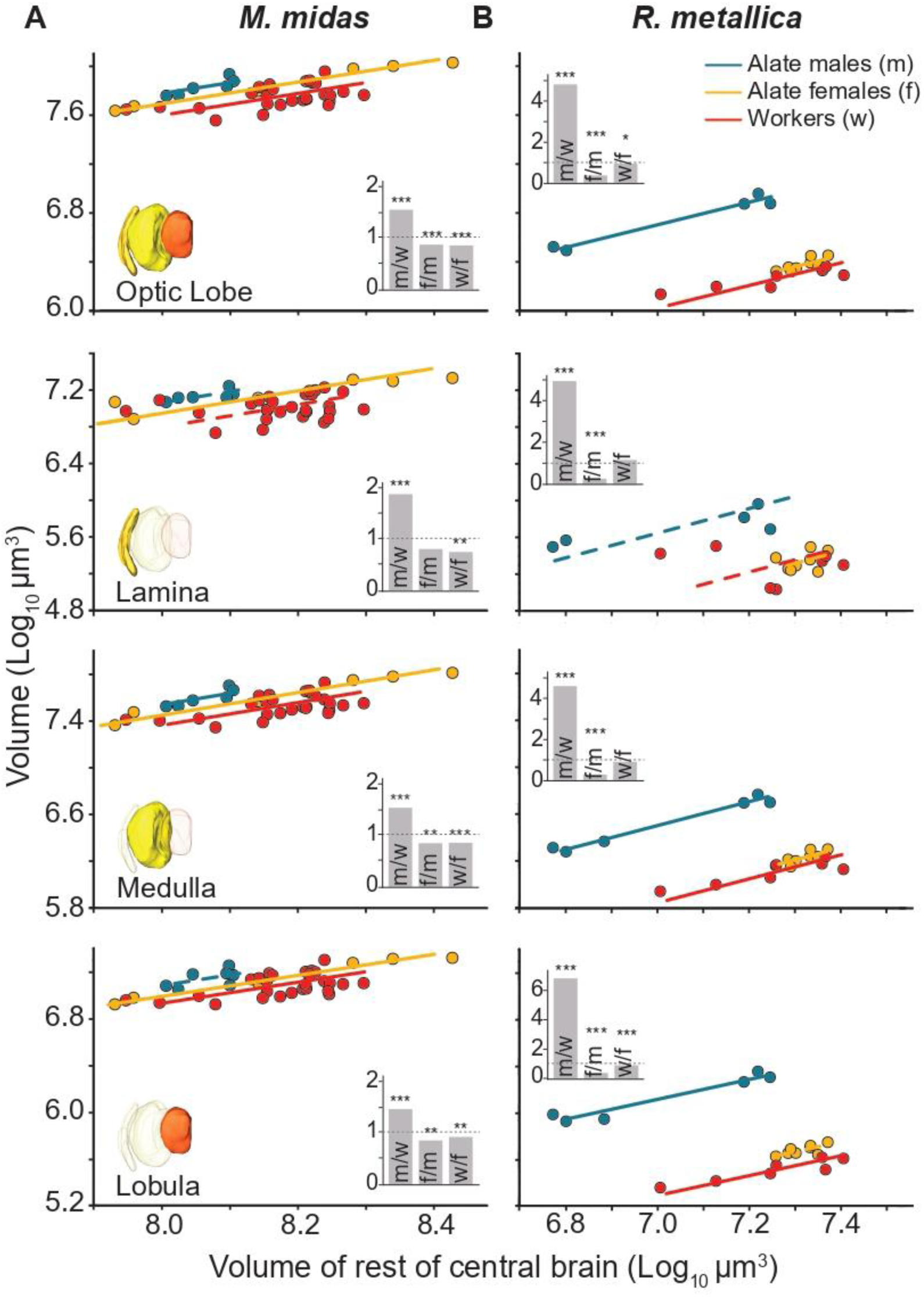
Scaling relationship of the volume of the optic lobes (OL) and the sub neuropils between the castes of *Myrmecia midas* and *Rytidoponera metallica*. Scaling relations of the whole optic lobe, lamina, medulla and lobula to the reference structure, the RoCB, for alate males (blue), alate females (yellow) and workers (red) of *M. midas* (A) and *R. metallica* (B). Each panel shows comparisons between castes of the same species. Regression lines are derived from SMA analysis and fitted to the data. Dashed lines represent non-significant linear regression models. The inset for each graph shows gsi for the region of interest. The gsi estimates the difference in the neuropil size between two castes of the same species and is calculated using the elevation of the regression line. From left to right, bars represent gsi calculated for alate males and workers (m/w), alate females and alate males (f/m) and workers and alate females (w/f). A gsi of 1 indicates that both castes have a similarly sized neuropils, shown by the horizontal dotted line within the bar graph. A gsi <1 indicates that the neuropil is larger in the second caste in the comparison and vice versa. If scaling slopes significantly differed between the three groups being compared, rendering it theoretically invalid to test for a true difference in volume/grade shift, no gsi was calculated (indicated by a missing inset). Stars indicate the significance level of difference in elevation between two regression lines, such that * p < 0.05, ** p < 0.01 and *** p < 0.001.

### Antennal Lobe (AL)

The ALs are the primary olfactory processing regions (Figure 1).

#### Comparison between species

Workers and males of *R. metallica* had larger ALs relative to the RoCB compared to their counterparts in *M. midas* (Figure 3A, 3C, second row panels). In alate females, AL volumes were relatively larger in *R. metallica*, however, they were not significantly bigger than the alate females of *M. midas* (Figure 3B, second row panel; Supplementary Table 2).

#### Comparison within species

The ALs showed caste-specific differences in volume only in *M. midas*. With respect to the RoCB volume, males had relatively larger ALs than workers. However, alate females did not differ significantly from males or workers (Figure 5A and inset within). In *R. metallica*, however, AL volumes did not vary significantly across castes (Figure 5B; Supplementary Table 3).

**Figure 5:**
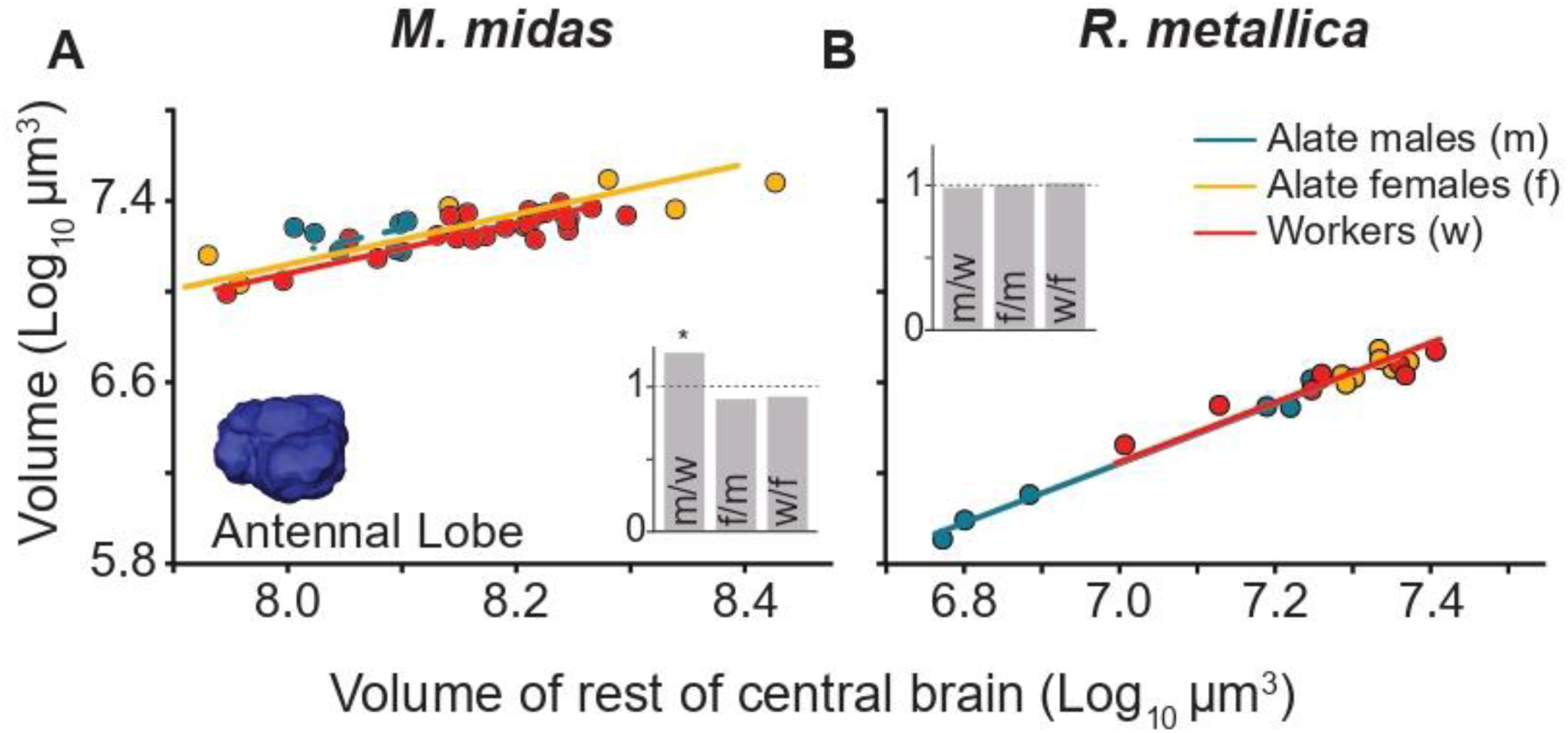
Scaling relationship of the volume of the Antennal Lobe (AL) between the castes of *Myrmecia midas* and *Rytidoponera metallica.* Scaling relations of the AL to the reference structure, the RoCB, for alate males (blue), alate females (yellow) and workers (red) of *M. midas* (A) and *R. metallica* (B). Figure conventions are as described in Figure 4.

### Mushroom Bodies (MB)

The MBs are the site of learning and memory in the insect brain. We divided the MBs into three sub-neuropils: the LIP, the COL, and the PED (which also includes the output regions of the MB; Figure 1).

#### Comparison between species

Within workers, relative to the RoCB, MB’s were significantly larger in *M. midas* than *R. metallica* (Figure 3A, third row panel). This difference was due to the significantly larger COL region (almost four times larger) in *M. midas*, while the LIP and PED were similar in size between the workers of the two species (Figure 3A, third row panel inset). Among alate females and males, the overall MB volume was comparable between species. In both castes, the LIP showed no notable differences and due to distinct scaling patterns, the COL could not be compared for absolute volume differences. Only in alate females, the PED was larger in *R. metallica* compared to *M. midas* (Figure 3B, third row panel inset; Supplementary Table 2).

#### Comparison within species

In both *M. midas* and *R. metallica*, relative to the RoCB, males consistently had the smallest MBs, alate females exhibited intermediate MB sizes, and workers possessed the largest. The magnitude of the volumetric difference was larger between alate female and males than between the two female castes (Figure 6A-B top panel and inset; Table 1). The general trend of MB size—with workers having the largest and males the smallest—was consistent across both the LIP and PED (Figure 6A and B, second and fourth row panels), with two exceptions: alate females and workers in both species shared similarly sized LIP regions (Figure 6A and B, second and fourth row panel insets). Scaling slopes of COL volumes among the three castes differed significantly in both species, rendering it theoretically invalid to test for a difference in volume or grade shift (Figure 6A-B third row panels; Supplementary Table 3).

**Figure 6:**
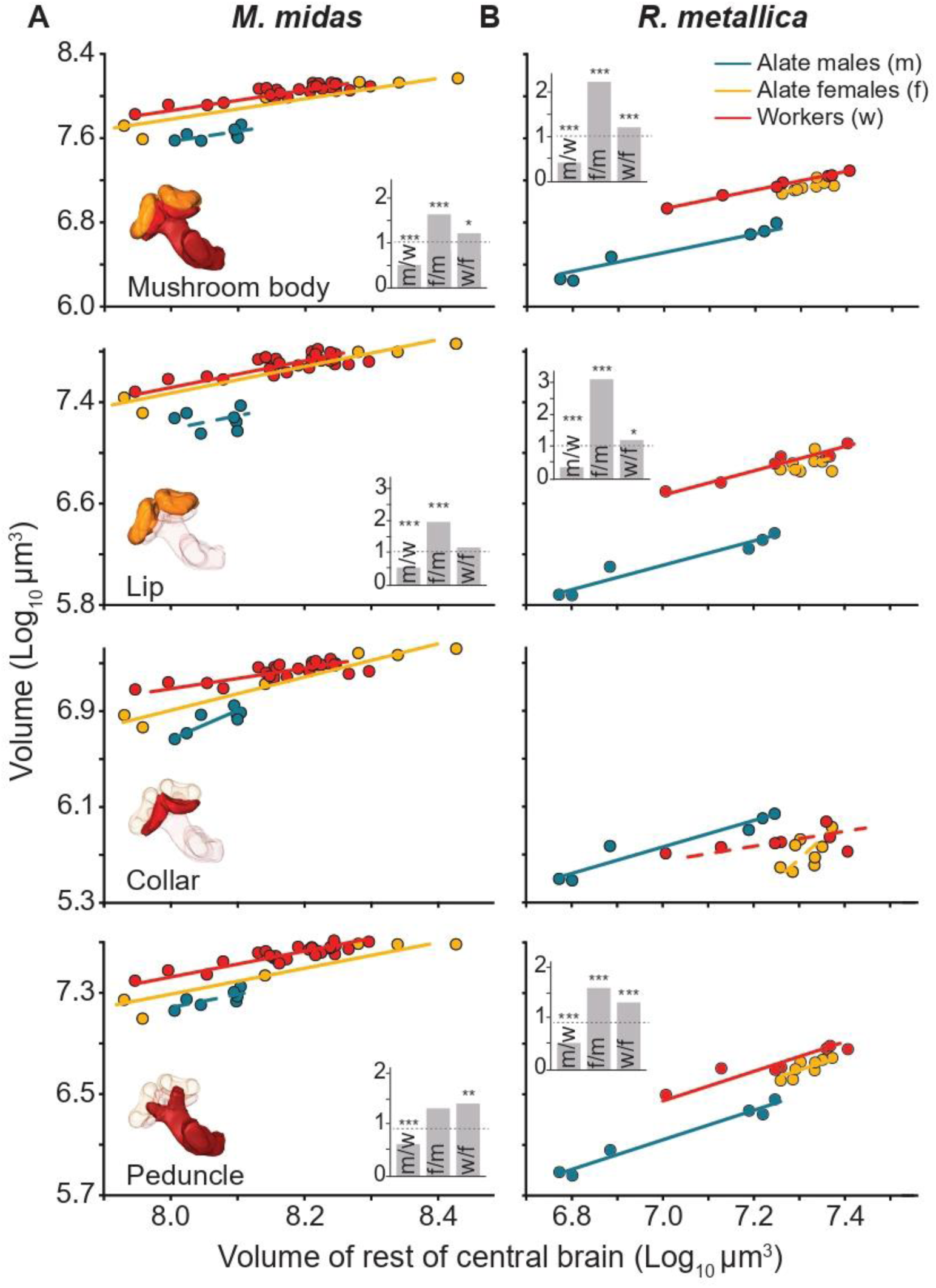
Scaling relationship of the volume of the Mushroom Body (MB) and the constituent neuropils between the castes of *Myrmecia midas* and *Rytidoponera metallica.* Scaling relations of the MB and its constituent neuropils to the reference structure, the RoCB, for alate males (blue), alate females (yellow) and workers (red) of *M. midas* (A) and *R. metallica* (B). Figure conventions are as described in Figure 4.

### Central Complex (CX)

The CX is a group of four neuropils: the FB, EB, PB and the paired NO (Figure 1).

#### Comparison between species

Workers of *R. metallica* showed significantly larger CXs relative to the RoCB, when compared to their *M. midas* counterparts (Figure 3A, bottom panel). This trend was seen in all sub-neuropils of the CX in the workers, except the PB, where scaling differed between the two species and thus could not be compared for grade shifts (Figure 3A, bottom panel inset). In alate females, the overall volumes of the CX, with the exception of the NO, were not significantly different (Figure 3B, bottom panel). The FB, EB and the PB showed varying scaling patterns in alate females (Figure 3B, bottom panel inset). Lastly, in males, the CX of *R. metallica* was significantly larger relative to the RoCB compared to *M. midas* (Figure 3C, bottom panel). This pattern was seen in all neuropils of the CX, except the NO, which scaled differently in both species (Figure 3C; Supplementary Table 2).

#### Comparison within species

In both species, relative to the RoCB, males had the largest CX (Figure 7A-B, top row panel). In *M. midas*, alate females had an intermediately sized CX, while workers had the smallest. Alate females of *M. midas* did not differ significantly from either males or workers (Figure 7A, top panel and inset). In *R. metallica*, no significant difference was found in the CX volumes among the two female castes; both were significantly smaller than the alate male (Figure 7B, top panel and inset).

**Figure 7:**
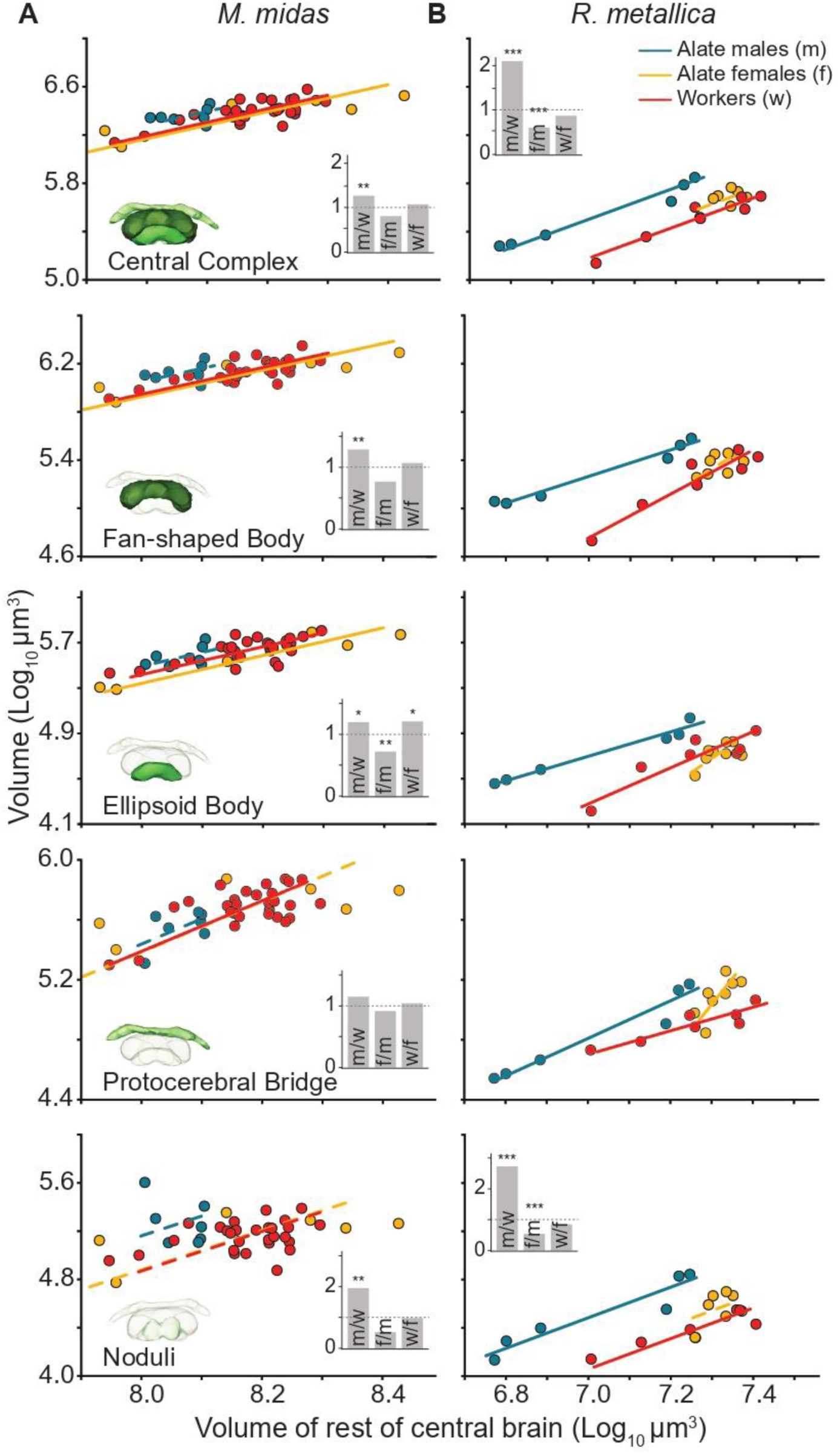
Scaling relationship of the volume of the Central Complex (CX) and the constituent neuropils between the castes of *Myrmecia midas* and *Rytidoponera metallica.* Scaling relations of the CX to the reference structure, the RoCB, for alate males (blue), alate females (yellow) and workers (red) of *M. midas* (A) and *R. metallica* (B). Figure conventions are as described in Figure 4.

The pattern seen in the CX of *M. midas*, with males having the largest, alate female intermediate, and workers with the smallest volumes relative to the RoCB, was also seen in the FB and the NO (Figure 7A, second and fifth row panel and insets). The EB, however, showed significant differences between all three castes, with the smallest volumes seen in alate females (Figure 7A, third row panel and inset). Lastly, there were no differences in the volume of the PB between the three castes in *M. midas* (Figure 7A, fourth row panel and inset). In *R. metallica*, the FB, EB, and the PB could not be compared for grade shifts or volumetric differences since the scaling slopes among the three castes were significantly different in each case (Figure 7B, second to fourth row panels and insets). In the NO, similar to the CX, we found no significant difference in the volume among the two female castes, both of which were significantly smaller than the alate male (Figure 7B, fifth row; Supplementary Table 3).

## Discussion

To investigate the relationship between lifestyles, ecologies and brain morphology, we measured neuropil volume across three castes in two species of Australian ants, *M. midas and R. metallica.* We identified variations in neuropil investment patterns throughout the brain, which likely reflect the considerable differences in lifestyle, mode of locomotion, visual ecology, and morphology that each species and caste exhibits. Our results also highlight areas where relative tissue investment appears to be conserved across castes or between species, pointing to shared evolutionary pressures or functional constraints that shape brain organisation in these species.

### Sensory processing centres

Our results indicate that investment in the OLs varies among different castes, reflecting the unique and varying visual demands faced by alate males, alate females, and workers. We observed significantly larger OL volumes in males when compared to both the female castes in the study species. This male-specific increase in investment in visual processing centres has been documented in several other ant species, including *Cataglyphis nodus* (Grob et al., 2021), *Atta vollenweideri* (Kübler et al., 2010), and *Camponotus japonicus* (Nishikawa et al., 2008).

Males and female alates must identify mating sites, with males relying heavily on their vision to intercept flying females mid-air while also fending off competing males. To achieve mating success while in flight, males require precise, fast, and accurate visual information. Males are known for their strong reliance on vision, evident from their disproportionately large compound eyes (Gronenberg and Hölldobler, 1999) and ocelli (Narendra et al., 2011). Male ants also have a higher number of ommatidia, the individual functional unit of a compound eye, compared to females of the same species (Gronenberg, 2008; Narendra et al., 2011). In some species, such as *Camponotus pennsylvanicus* and *Camponotus consobrinus*, both male and female alates have more ommatidia and larger eyes than workers of comparable head size (Klotz et al., 1992; Narendra et al., 2016). A greater number of ommatidia, along with larger lenses and smaller interommatidial angles, increases the spatial resolution of the eye and thus shows the importance of vision in both reproductive castes (Barlow, 1952; Jander and Jander, 2002; Kelber et al., 2006; Narendra et al., 2011; Somanathan et al., 2009; Spaethe and Chittka, 2003).

The mode of locomotion may also play a role in defining the investment in the OLs. Male alates spend a significant fraction of their lives on the wing, while female alates are also capable of flight. Workers, on the other hand, are purely ambulatory. Flying may thus necessitate better visual capabilities, such as higher temporal resolution, to allow males and females to locate mates successfully. Other insects that face similar visual processing challenges due to their aerial lifestyles also demonstrate well-developed visual processing centres. For instance, large and complex OLs can also be found in dragonflies, which must quickly respond to small, fast-moving targets while hunting (Fabian et al., 2020).

Our study revealed significant differences in AL volumes between males and workers of *M. midas*. Males possessed larger ALs, a pattern also observed in other ants such as *C. nodus* (Grob et al., 2021), *C. japonicus* (Nishikawa et al., 2008), *C. sericeus and C. compressus* (Mysore et al., 2009). Males in several insect species, including ants (Nishikawa et al., 2008; Kübler et al., 2010), honey bees (Arnold et al., 1985), and moths (Hansson et al., 1991; Zhao et al., 2016), possess enlarged glomeruli (the distinct processing units in the AL) specialised for detecting sex pheromones. While sex pheromones likely influence the mating behaviour of *M. midas*, this remains to be confirmed. Furthermore, no differences were detected in the AL volumes between the two female castes of *M. midas*. This pattern contrasts with that of honey bees, where queens have slightly smaller AL volumes compared to workers, resulting in reduced performance in olfactory learning tasks (Groh and Rössler, 2008). Unlike honey bee queens, alate females of *M. midas* engage in foraging and nest maintenance during the establishment phase of their colony (Reid et al., 2013; Haskins and Haskins, 1950), necessitating a sense of smell comparable to that of workers, who subsequently assume these duties.

In contrast to *M. midas*, there were no significant differences in AL volume in the three castes of *R. metallica*. Additionally, the ALs in all castes of *R. metallica* were consistently larger relative to RoCB when compared to their counterparts in *M. midas*, suggesting a greater reliance on olfactory cues or perhaps a less complex cuticular hydrocarbon profile (Marty et al., 2025). Similarly, diurnal and nocturnal dung beetles that strongly rely on olfactory cues for many key behaviours, such as finding specific types of dung to feed on, also do not show differences in ALs, despite differences in behaviour (Immonen et al., 2017).

Generally, a trade-off is noted between the visual and olfactory sensory regions— nocturnal insects invest more heavily in olfactory processing regions, while diurnal insects invest more in visual processing regions (bull ants: Sheehan et al. 2019; hawk moths: Stöckl et al. 2016; paper wasps: O’Donnell et al. 2013), thus taking advantage of the readily available cues in their surroundings. Surprisingly, our study reveals an inverse pattern: nocturnal *M. midas* workers had larger visual processing regions than diurnal *R. metallica* workers, while the latter exhibited larger olfactory processing regions.

Larger OLs and more facets in the compound eye of the nocturnal *M. midas* workers may be an adaptation to improve vision in dim-light conditions. Workers of *M. midas* have a head width of 3.91 ± 0.11 mm, possess approximately 3,590 ± 88 facets in each compound eye (Ogawa et al., 2019). Workers of *R. metallica* are significantly smaller, with a head width of 1.44 ± 0.07 mm and an average of 275 ± 12 facets in their eyes (Palavalli-Nettimi and Narendra, 2018). Thus, relative to their head size, *M. midas* workers have approximately five times more facets than *R. metallica* workers, which suggests they have a significantly higher sampling resolution. Workers of *M. midas* also have larger lenses compared to *R. metallica* (31.62 µm vs 18.5 µm), which is a visual adaptation to enhance the light gathering capacity of the eye, essential for dim light conditions (Ogawa et al., 2019, Palavalli-Nettimi and Narendra, 2018, Narendra et al., 2011).

### Higher-order integration

In insect brains, MBs are centres of multimodal associative learning and play a fundamental role in visual navigation (Buehlmann et al., 2020; Kahmi et al., 2020; Mizunami et al., 1998; Modi et al., 2020; Strausfeld et al., 1998). In our study, males of both species possessed the smallest MBs. Since males perish shortly after copulation (Hölldobler and Wilson, 1990), they do not need to acquire visual memories, such as those used to locate a nest after a foraging trip, leading to lower investment in memory centres (Ehmer and Gronenberg, 2004), a pattern also seen in termites (Merchant and Zhou, 2024) and other ants, such as *C. nodus* (Grob et al., 2021).

In contrast, workers of both *M. midas* and *R. metallica* possess the largest MBs among the three castes. Workers perform diverse tasks requiring significant behavioural flexibility, particularly in the domain of associative memory. For example, *M. midas* foragers learn and memorise visual panoramas and landmarks to locate their nest following a foraging trip (Reid et al., 2011). Similarly, young queens of both *R. metallica* (Ward, 1986) and *M. midas* (Reid et al., 2013; Haskins and Haskins, 1950) also face navigational challenges for a brief period of their lives while establishing new nests. Visual homing requires better processing capabilities in brain areas involved with visual image recognition, which likely contributes to the observed differences in MB volume between castes.

Consistent with this, the MB calyx in ants receives segregated sensory input, with the LIP dominated by olfactory projections and the COL by visual projections. In both species, workers and queens possess significantly larger LIP regions, suggesting the two female castes invest most in olfactory associative memory. The underlying reason for the volumetric differences in the calyces between castes may be similar to that in *C. nodus*: workers and queens possess nearly double the number of microglomeruli in the MB calyces compared to males (Grob et al., 2021). While statistical comparisons of COL size between castes were not possible, workers of *M. midas* had a significantly larger relative COL than *R. metallica*, that reflects the strong reliance of vision in *Myrmecia* ants (e.g., Reid et al., 2013; Kamhi et al., 2020).

The insect CX is involved in many functions, including modulating internal states (such as sleep), transforming sensory signals into internal representations of head direction, encoding travel direction, and initiating steering commands (reviewed in Heinze, 2023). Across insects, the CX is highly morphologically conserved (Honkanen et al., 2019; de Vries et al., 2021; Farnworth et al., 2025). Males of both species studied here possessed the largest CXs, which aligns with previous observations in other ant species (Grob et al., 2021; Kübler et al., 2010; Nishikawa et al., 2008). Of the four CX neuropils, the greatest difference was observed in the noduli. The noduli are innervated by neurons that encode rotational and translational speed, as well as neurons that project to the FB and are involved in encoding travel direction (Lyu et al., 2022; Lu et al., 2022; Stone et al., 2017). Enlarged noduli may be attributable to the fact that males spend most of their active life in flight and therefore rely more on accurately monitoring their flight speed and travel direction. Additionally, male ants need to target fast-flying objects in a cluttered background while competing with other males. This increased investment in the CX has been suggested to ‘promote efficacy in male mating behaviour’ among other advantages (Grob et al., 2021).

## Conclusion

In summary, we found clear differences in the relative volumes of functionally distinct brain regions both between species and among castes within the same species. Adaptations in each caste appear to closely match their specific life histories, tasks, modes of locomotion, and the light environments they occupy.

## Acknowledgements

We acknowledge the Traditional Custodians of the land where we carried out our research, the Wallumattagal Clan of the Dharug Nation. We thank Sue Lindsay and Arthur Chien, for providing us access to microscopy facilities. The research was supported by an International Research Training Program Stipend Scholarship to SE, and an Australian Research Council Discovery Project grant DP220102836 to AN.

## Author contributions

Conceptualization: AN. Methodology: SE, ZBVS, MES, AN. Formal analysis: SE, MES, AN. Investigation: SE, ZBVS, MES, AN. Resources: FP, AN. Data curation: SE, AN. Writing, original draft: SE. Writing, review & editing: SE, MES, FP, AN. Visualization: SE, MES, AN. Funding acquisition: AN.

## References

Arnold, G., Masson, C., & Budharugsa, S. (1985). Comparative study of the antennal lobes and their afferent pathway in the worker bee and the drone (*Apis mellifera*). Cell & Tissue Research, 242(3), 593–605.

Barlow, H. B. (1952). The size of ommatidia in apposition eyes. Journal of Experimental Biology, 29(4), 667–674.

Bouchebti, S., & Arganda, S. (2020). Insect lifestyle and evolution of brain morphology. Current Opinion in Insect Science, 42, 90–96.

Buehlmann, C., Wozniak, B., Goulard, R., Webb, B., Graham, P., & Niven, J. E. (2020). Mushroom bodies are required for learned visual navigation, but not for innate visual behavior, in ants. Current Biology, 30(17), 3438–3443.

Cabirol, A., Cope, A. J., Barron, A. B., & Devaud, J. M. (2018). Relationship between brain plasticity, learning and foraging performance in honey bees. PLoS One, 13(4), e0196749.

Collett, T., Graham, P., & Heinze, S. (2025). The neuroethology of ant navigation. Current Biology, 35(3), R110–R124.

Couto, A., Young, F. J., Atzeni, D., Marty, S., Melo-Flórez, L., Hebberecht, L., Monllor, M., Neal, C., Cicconardi, F., McMillan, W. O., et al. (2023). Rapid expansion and visual specialisation of learning and memory centres in the brains of *Heliconiini* butterflies. Nature Communications, 14(1), 4024.

Diel, E. E., Lichtman, J. W., & Richardson, D. S. (2020). Tutorial: avoiding and correcting sample-induced spherical aberration artifacts in 3D fluorescence microscopy. Nature Protocols, 15(9), 2773–2784.

Ehmer, B., & Gronenberg, W. (2004). Mushroom body volumes and visual interneurons in ants: comparison between sexes and castes. Journal of Comparative Neurology, 469(2), 198–213.

Fabian, J. M., el Jundi, B., Wiederman, S. D., & O’Carroll, D. C. (2020). The complex optic lobe of dragonflies. bioRxiv, 2020–05.

Farnworth, M. S., Loupasaki, T., Couto, A., & Montgomery, S. H. (2024). Mosaic evolution of a learning and memory circuit in *Heliconiini* butterflies. Current Biology, 34(22), 5252–5262.

Farnworth, M. S., Toh, Y. P., Loupasaki, T., Hodge, E. A., el Jundi, B., & Montgomery, S. H. (2025). Distinct evolutionary trajectories of two integration centres, the central complex and mushroom bodies, across *Heliconiini* butterflies. bioRxiv, 2025–05.

Farris, S. M., Robinson, G. E., & Fahrbach, S. E. (2001). Experience- and age-related outgrowth of intrinsic neurons in the mushroom bodies of the adult worker honeybee. Journal of Neuroscience, 21(16), 6395–6404.

Freas, C. A., Narendra, A., Murray, T., & Cheng, K. (2024). Polarised moonlight guides nocturnal bull ants home. eLife, 13, RP97615.

Ghaffar, H., Larsen, J. R., Booth, G. M., & Perkes, R. (1984). General morphology of the brain of the blind cave beetle, *Neaphaenops tellkampfii* Erichson (Coleoptera: Carabidae). International Journal of Insect Morphology and Embryology, 13(5-6), 357–371.

Greiner, B., Narendra, A., Reid, S. F., Dacke, M., Ribi, W. A., & Zeil, J. (2007). Eye structure correlates with distinct foraging-bout timing in primitive ants. Current Biology, 17(20), R879–R880.

Grob, R., Fleischmann, P. N., Grübel, K., Wehner, R., & Rössler, W. (2017). The role of celestial compass information in *Cataglyphis* ants during learning walks and for neuroplasticity in the central complex and mushroom bodies. Frontiers in Behavioral Neuroscience, 11, 226.

Grob, R., Fleischmann, P. N., & Rössler, W. (2019). Learning to navigate–how desert ants calibrate their compass systems. Neuroforum, 25(2), 109–120.

Grob, R., Heinig, N., Grübel, K., Rössler, W., & Fleischmann, P. N. (2021). Sex-specific and caste-specific brain adaptations related to spatial orientation in *Cataglyphis* ants. Journal of Comparative Neurology, 529(18), 3882–3892.

Groh, C., & Rössler, W. (2008). Caste-specific postembryonic development of primary and secondary olfactory centers in the female honeybee brain. Arthropod Structure & Development, 37(6), 459–468.

Groh, C., & Rössler, W. (2020). Analysis of synaptic microcircuits in the mushroom bodies of the honeybee. Insects, 11(1), 43.

Gronenberg, W. (2008). Structure and function of ant (Hymenoptera: Formicidae) brains: strength in numbers. Myrmecological News, 11, 25–36.

Gronenberg, W., Heeren, S., & Hölldobler, B. (1996). Age-dependent and task-related morphological changes in the brain and the mushroom bodies of the ant *Camponotus floridanus*. Journal of Experimental Biology, 199(9), 2011–2019.

Gronenberg, W., & Hölldobler, B. (1999). Morphologic representation of visual and antennal information in the ant brain. Journal of Comparative Neurology, 412(2), 229–240.

Hansson, B. S., Christensen, T. A., & Hildebrand, J. G. (1991). Functionally distinct subdivisions of the macroglomerular complex in the antennal lobe of the male sphinx moth *Manduca sexta*. Journal of Comparative Neurology, 312(2), 264–278.

Haskins, C. P., & Haskins, E. F. (1950). Notes on the biology and social behavior of the archaic ponerine ants of the genera *Myrmecia* and *Promyrmecia*. Annals of the Entomological Society of America, 43(4), 461–491.

Heinze, S. (2023). The insect central complex. In Oxford Research Encyclopedia of Neuroscience. Oxford University Press.

Hölldobler, B., & Wilson, E. O. (1990). The Ants. Harvard University Press.

Honkanen, A., Adden, A., da Silva Freitas, J., & Heinze, S. (2019). The insect central complex and the neural basis of navigational strategies. Journal of Experimental Biology, 222(Suppl 1), jeb188854.

Hughes, L. (1991). The relocation of ant nest entrances: Potential consequences for ant-dispersed seeds. Australian Journal of Ecology, 16(2), 207–214.

Immonen, E. V., Dacke, M., Heinze, S., & el Jundi, B. (2017). Anatomical organization of the brain of a diurnal and a nocturnal dung beetle. Journal of Comparative Neurology, 525(8), 1879–1908.

Jander, U., & Jander, R. (2002). Allometry and resolution of bee eyes (Apoidea). Arthropod Structure & Development, 30(3), 179–193.

Joseph, A. (2023). Visual navigation in ants while walking backwards (Master’s thesis). Macquarie University.

Kamhi, J. F., Barron, A. B., & Narendra, A. (2020). Vertical lobes of the mushroom bodies are essential for view-based navigation in Australian *Myrmecia* ants. Current Biology, 30(17), 3432–3437.

Kelber, A., Warrant, E. J., Pfaff, M., Wallén, R., Theobald, J. C., Wcislo, W. T., & Raguso, R. A. (2006). Light intensity limits foraging activity in nocturnal and crepuscular bees. Behavioral Ecology, 17(1), 63–72.

Klotz, J., Reid, B., & Gordon, W. (1992). Variation of ommatidia number as a function of worker size in *Camponotus pennsylvanicus* (DeGeer) (Hymenoptera: Formicidae). Insectes Sociaux, 39(2), 233–236.

Kübler, L. S., Kelber, C., & Kleineidam, C. J. (2010). Distinct antennal lobe phenotypes in the leaf-cutting ant (*Atta vollenweideri*). Journal of Comparative Neurology, 518(3), 352–365.

Kühn-Bühlmann, S., & Wehner, R. (2006). Age-dependent and task-related volume changes in the mushroom bodies of visually guided desert ants, *Cataglyphis bicolor*. Journal of Neurobiology, 66(6), 511–521.

Lösel, P. D., van de Kamp, T., Jayme, A., Ershov, A., Faragó, T., Pichler, O., Tan, J. N., Aadepu, N., Bremer, S., Chilingaryan, S. A., et al. (2020). Introducing Biomedisa as an open-source online platform for biomedical image segmentation. Nature Communications, 11(1), 5577.

Lu, J., Behbahani, A. H., Hamburg, L., Westeinde, E. A., Dawson, P. M., Lyu, C., Maimon, G., Dickinson, M. H., Druckmann, S., & Wilson, R. I. (2022). Transforming representations of movement from body-to world-centric space. Nature, 601(7891), 98–104.

Lyu, C., Abbott, L., & Maimon, G. (2022). Building an allocentric travelling direction signal via vector computation. Nature, 601(7891), 92–97.

Marty, S., Couto, A., Dawson, E. H., Brard, N., d’Ettorre, P., Montgomery, S. H., & Sandoz, J. C. (2025). Ancestral complexity and constrained diversification of the ant olfactory system. Proceedings of the Royal Society B, 292(2045), 20250662.

Merchant, A., & Zhou, X. (2024). Caste-biased patterns of brain investment in the subterranean termite *Reticulitermes flavipes*. iScience, 27(6).

Mizunami, M., Weibrecht, J. M., & Strausfeld, N. J. (1998). Mushroom bodies of the cockroach: their participation in place memory. Journal of Comparative Neurology, 402(4), 520–537.

Modi, M. N., Shuai, Y., & Turner, G. C. (2020). The *Drosophila* mushroom body: from architecture to algorithm in a learning circuit. Annual Review of Neuroscience, 43(1), 465–484.

Moser, J. C., Reeve, J. D., Bento, J. M. S., Della Lucia, T. M., Cameron, R. S., & Heck, N. M. (2004). Eye size and behaviour of day-and night-flying leafcutting ant alates. Journal of Zoology, 264(1), 69–75.

Mysore, K., Subramanian, K., Sarasij, R., Suresh, A., Shyamala, B. V., VijayRaghavan, K., & Rodrigues, V. (2009). Caste and sex specific olfactory glomerular organization and brain architecture in two sympatric ant species *Camponotus sericeus* and *Camponotus compressus* (Fabricius, 1798). Arthropod Structure & Development, 38(6), 485–497.

Nakanishi, A., Nishino, H., Watanabe, H., Yokohari, F., & Nishikawa, M. (2009). Sex-specific antennal sensory system in the ant *Camponotus japonicus*: structure and distribution of sensilla on the flagellum. Cell and Tissue Research, 338(1), 79–97.

Narasimhan, S., Villar, M. E., Chiara, V., Arganda-Carreras, I., Arganda, S., Witek, M., & Sanmartín-Villar, I. (2025). Complexity shapes uniqueness: Neuropil volumes and synaptic clusters shape behavioural plasticity under challenging environments in the invasive Argentine ants. bioRxiv, 2025–03.

Narendra, A., Kamhi, J. F., & Ogawa, Y. (2017). Moving in dim light: behavioral and visual adaptations in nocturnal ants. Integrative and Comparative Biology, 57(5), 1104–1116.

Narendra, A., Ramirez-Esquivel, F., & Ribi, W. A. (2016). Compound eye and ocellar structure for walking and flying modes of locomotion in the Australian ant, *Camponotus consobrinus*. Scientific Reports, 6(1), 22331.

Narendra, A., Reid, S. F., Greiner, B., Peters, R. A., Hemmi, J. M., Ribi, W. A., & Zeil, J. (2011). Caste-specific visual adaptations to distinct daily activity schedules in Australian *Myrmecia* ants. Proceedings of the Royal Society B, 278(1709), 1141–1149.

Narendra, A., & Ribi, W. A. (2017). Ocellar structure is driven by the mode of locomotion and activity time in *Myrmecia* ants. Journal of Experimental Biology, 220(23), 4383–4390.

Nishikawa, M., Nishino, H., Misaka, Y., Kubota, M., Tsuji, E., Satoji, Y., Ozaki, M., & Yokohari, F. (2008). Sexual dimorphism in the antennal lobe of the ant *Camponotus japonicus*. Zoological Science, 25(2), 195–204.

O’Donnell, S., Clifford, M. R., DeLeon, S., Papa, C., Zahedi, N., & Bulova, S. J. (2013). Brain size and visual environment predict species differences in paper wasp sensory processing brain regions (Hymenoptera: Vespidae, Polistinae). Brain, Behavior and Evolution, 82(3), 177–184.

Ogawa, Y., Ryan, L. A., Palavalli-Nettimi, R., Seeger, O., Hart, N. S., & Narendra, A. (2019). Spatial resolving power and contrast sensitivity are adapted for ambient light conditions in Australian *Myrmecia* ants. Frontiers in Ecology and Evolution, 7, 18.

Ott, S. R., & Rogers, S. M. (2010). Gregarious desert locusts have substantially larger brains with altered proportions compared with the solitarious phase. Proceedings of the Royal Society B, 277(1697), 3087–3096.

Palavalli-Nettimi, R., & Narendra, A. (2018). Miniaturisation decreases visual navigational competence in ants. Journal of Experimental Biology, 221(7), jeb177238.

Penmetcha, B., Ogawa, Y., Ribi, W. A., & Narendra, A. (2019). Ocellar structure of African and Australian desert ants. Journal of Comparative Physiology A, 205(5), 699–706.

Reid, S. F., Narendra, A., Hemmi, J. M., & Zeil, J. (2011). Polarised skylight and the landmark panorama provide night-active bull ants with compass information during route following. Journal of Experimental Biology, 214(3), 363–370.

Reid, S. F., Narendra, A., Taylor, R. W., & Zeil, J. (2013). Foraging ecology of the night-active bull ant *Myrmecia pyriformis*. Australian Journal of Zoology, 61(2), 170–177.

Riveros, A. J., & Gronenberg, W. (2010). Brain allometry and neural plasticity in the bumblebee *Bombus occidentalis*. Brain, Behavior and Evolution, 75(2), 138–148.

Rozanski, A. N., Cini, A., Lopreto, T. E., Gandia, K. M., Hauber, M. E., Cervo, R., & Uy, F. M. (2022). Differential investment in visual and olfactory brain regions is linked to the sensory needs of a wasp social parasite and its host. Journal of Comparative Neurology, 530(4), 756–767.

Scholl, C., Wang, Y., Krischke, M., Mueller, M. J., Amdam, G. V., & Rössler, W. (2014). Light exposure leads to reorganization of microglomeruli in the mushroom bodies and influences juvenile hormone levels in the honeybee. Developmental Neurobiology, 74(11), 1141–1153.

Seid, M. A., & Wehner, R. (2009). Delayed axonal pruning in the ant brain: a study of developmental trajectories. Developmental Neurobiology, 69(6), 350–364.

Sheehan, Z. B., Kamhi, J. F., Seid, M. A., & Narendra, A. (2019). Differential investment in brain regions for a diurnal and nocturnal lifestyle in Australian *Myrmecia* ants. Journal of Comparative Neurology, 527(7), 1261–1277.

Somanathan, H., Kelber, A., Borges, R. M., Wallén, R., & Warrant, E. J. (2009). Visual ecology of Indian carpenter bees II: adaptations of eyes and ocelli to nocturnal and diurnal lifestyles. Journal of Comparative Physiology A, 195(6), 571–583.

Spaethe, J., & Chittka, L. (2003). Interindividual variation of eye optics and single object resolution in bumblebees. Journal of Experimental Biology, 206(19), 3447–3453.

Stieb, S. M., Muenz, T. S., Wehner, R., & Rössler, W. (2010). Visual experience and age affect synaptic organization in the mushroom bodies of the desert ant *Cataglyphis fortis*. Developmental Neurobiology, 70(6), 408–423.

Stöckl, A., Heinze, S., Charalabidis, A., el Jundi, B., Warrant, E., & Kelber, A. (2016). Differential investment in visual and olfactory brain areas reflects behavioural choices in hawk moths. Scientific Reports, 6(1), 26041.

Stone, T., Webb, B., Adden, A., Weddig, N. B., Honkanen, A., Templin, R., Wcislo, W., Scimeca, L., Warrant, E., & Heinze, S. (2017). An anatomically constrained model for path integration in the bee brain. Current Biology, 27(20), 3069–3085.

Strausfeld, N. J., Hansen, L., Li, Y., Gomez, R. S., & Ito, K. (1998). Evolution, discovery, and interpretations of arthropod mushroom bodies. Learning & Memory, 5(1), 11–37.

Thomas, M. (2002). Nest site selection and longevity in the ponerine ant *Rhytidoponera metallica* (Hymenoptera, Formicidae). Insectes Sociaux, 49, 147–152.

de Vries, L., Pfeiffer, K., Trebels, B., Adden, A. K., Green, K., Warrant, E., & Heinze, S. (2017). Comparison of navigation-related brain regions in migratory versus non-migratory noctuid moths. Frontiers in Behavioral Neuroscience, 11, 158.

Ward, P. S. (1986). Functional queens in the Australian greenhead ant, *Rhytidoponera metallica* (Hymenoptera: Formicidae). Psyche: A Journal of Entomology, 93(1-2), 1–12.

Warton, D. I., Duursma, R. A., Falster, D. S., & Taskinen, S., et al. (2012). smatr 3–an R package for estimation and inference about allometric lines. Methods in Ecology and Evolution, 3(2), 257–259.

Withers, G. S., Fahrbach, S. E., & Robinson, G. E. (1993). Selective neuroanatomical plasticity and division of labour in the honeybee. Nature, 364(6434), 238–240.

Yilmaz, A., Lindenberg, A., Albert, S., Grübel, K., Spaethe, J., Rössler, W., & Groh, C. (2016). Age-related and light-induced plasticity in opsin gene expression and in primary and secondary visual centers of the nectar-feeding ant *Camponotus rufipes*. Developmental Neurobiology, 76(9), 1041–1057.

Zhao, X. C., Ma, B. W., Berg, B. G., Xie, G. Y., Tang, Q. B., & Guo, X. R. (2016). A global-wide search for sexual dimorphism of glomeruli in the antennal lobe of female and male *Helicoverpa armigera*. Scientific Reports, 6, 35204.

