## Supplementary material for "Caste- and Sex-Specific Differential Investment in Brain Regions of Australian Ants": Table S1

Supplementary Table 1: Absolute volume of neuropils and headwidth of workers, alate males (males) and alate females (females) of *Myrmecia midas* and *Rhytidoponera metallica.*

| **Species** | **Caste** | **LA (µm^3^)** | **ME (µm^3^)** | **LO (µm^3^)** | **OL (µm^3^)** | **AL (µm^3^)** |
| --- | --- | --- | --- | --- | --- | --- |
| *M. midas* | Worker | 9614885 | 40041244 | 13702118 | 63358247 | 18972754 |
| *M. midas* | Worker | 7706247.5 | 36576804 | 13361707 | 57644758.5 | 18099158 |
| *M. midas* | Worker | 10593422 | 31168596 | 10367414 | 52129432 | 18671144 |
| *M. midas* | Worker | 5519153 | 22588708 | 8573276 | 36681137 | 13993953 |
| *M. midas* | Worker | 9740461 | 35673904 | 12783525 | 58197890 | 21674984 |
| *M. midas* | Worker | 9362312 | 26084782 | 9243072 | 44690166 | 9816630 |
| *M. midas* | Worker | 9687637 | 28903832 | 10969934 | 49561403 | 18276080 |
| *M. midas* | Worker | 8200183.5 | 33304248 | 11447070 | 52951501.5 | 19422548 |
| *M. midas* | Worker | 7120490 | 38373332 | 13278900 | 58772722 | 22608736 |
| *M. midas* | Worker | 9162073 | 29513396 | 9931740 | 48607209 | 17593808 |
| *M. midas* | Worker | 9230463 | 33235956 | 11534556 | 54000975 | 21484006 |
| *M. midas* | Worker | 12237120 | 25683526 | 8845994 | 46766640 | 11171904 |
| *M. midas* | Worker | 11239199 | 35237728 | 13731821 | 60208748 | 17699796 |
| *M. midas* | Worker | 13239193 | 41798816 | 15421471 | 70459480 | 22244490 |
| *M. midas* | Worker | 12119717 | 41072420 | 14039736 | 67231873 | 21605508 |
| *M. midas* | Worker | 14186893 | 44811832 | 15846791 | 74845516 | 22821704 |
| *M. midas* | Worker | 10307851 | 31621158 | 10638086 | 52567095 | 19181328 |
| *M. midas* | Worker | 14020944 | 42832148 | 16166565 | 73019657 | 21982634 |
| *M. midas* | Worker | 5951604.5 | 24897872 | 9647949 | 40497425.5 | 17194548 |
| *M. midas* | Worker | 15524139 | 40898520 | 15746275 | 72168934 | 22248172 |
| *M. midas* | Worker | 9471543 | 32574110 | 12493360 | 54539013 | 19607628 |
| *M. midas* | Worker | 9259065 | 35621368 | 12814967 | 57695400 | 21035046 |
| *M. midas* | Worker | 11798701 | 38225968 | 14787765 | 64812434 | 17018556 |
| *M. midas* | Worker | 15287500 | 45343972 | 15147843 | 75779315 | 17025918 |
| *M. midas* | Worker | 16629806 | 52901124 | 19826480 | 89357410 | 24601426 |
| *M. midas* | Worker | 9022599 | 26719512 | 10044615 | 45786726 | 17125618 |
| *M. midas* | Worker | 14959209 | 34249252 | 12536751 | 61745212 | 23400060 |
| *M. midas* | Worker | 7817601.5 | 29654404 | 11011708 | 48483713.5 | 20439378 |
| *M. midas* | Male | 11798449 | 33880112 | 12193462 | 57872023 | 19163636 |
| *M. midas* | Male | 13091601 | 34106568 | 11413834 | 58612003 | 18082166 |
| *M. midas* | Male | 17398868 | 50248728 | 17922500 | 85570096 | 19926208 |

| **LIP (µm^3^)** | **COL (µm^3^)** | **PED (µm^3^)** | **MB (µm^3^)** |
| --- | --- | --- | --- |
| 50127912 | 18525993 | 37480872 | 106134777 |
| 45861116 | 14641451.5 | 38355200 | 98857767.5 |
| 50118046 | 19155624 | 40349584 | 109623254 |
| 38023172 | 12309734 | 35318244 | 85651150 |
| 52572242 | 17096371.5 | 50583744 | 120252357.5 |
| 30420865 | 12057409.5 | 24892212 | 67370486.5 |
| 40791166 | 15180680 | 38495040 | 94466886 |
| 47040592 | 17074141 | 43760468 | 107875201 |
| 53166788 | 19617170 | 44756160 | 117540118 |
| 43243382 | 15359750.5 | 36819476 | 95422608.5 |
| 58220950 | 19328405 | 45202380 | 122751735 |
| 38268866 | 13819918 | 30011858 | 82100642 |
| 54681074 | 18438718 | 41413388 | 114533180 |
| 55280034 | 17961760 | 38870096 | 112111890 |
| 56895784 | 16615494 | 42539224 | 116050502 |
| 63120914 | 20301965 | 45687728 | 129110607 |
| 48974550 | 17982064 | 46076212 | 113032826 |
| 65909304 | 19286884 | 42691304 | 127887492 |
| 46082880 | 15570399.5 | 39287460 | 100940739.5 |
| 53579702 | 19363265 | 41440156 | 114383123 |
| 54943718 | 19182052 | 45001412 | 119127182 |
| 60229822 | 19831244 | 47117988 | 127179054 |
| 51256740 | 19374935 | 34136788 | 104768463 |
| 60386278 | 20811081 | 39591036 | 120788395 |
| 62193642 | 21647912 | 45841988 | 129683542 |
| 40039722 | 13641033 | 27851302 | 81532057 |
| 50008522 | 16295169 | 44303924 | 110607615 |
| 54958670 | 19479914 | 51636216 | 126074800 |
| 18807252 | 4644628.5 | 14418135 | 37870015.5 |
| 20616406 | 5196952 | 17741112 | 43554470 |
| 17571412 | 7521137 | 17096686 | 42189235 |

| **Species** | **Caste** | **LA (µm^3^)** | **ME (µm^3^)** | **LO (µm^3^)** | **OL (µm^3^)** | **AL (µm^3^)** |
| --- | --- | --- | --- | --- | --- | --- |
| *M. midas* | Male | 14295032 | 46099084 | 14947791 | 75341907 | 20471778 |
| *M. midas* | Male | NA | NA | 12323535 | NA | 15068933 |
| *M. midas* | Male | 13254654 | 40039396 | 15457878 | 68751928 | 15335101 |
| *M. midas* | Male | 13283926 | 37896576 | 15167513 | 66348015 | 14995433 |
| *M. midas* | Female | 19550988 | 59509956 | 20292936 | 99353880 | 23085988 |
| *M. midas* | Female | 20270060 | 55611000 | 18775374 | 94656434 | 31256764 |
| *M. midas* | Female | 21096624 | 63845516 | 20659760 | 105601900 | 30251724 |
| *M. midas* | Female | 7797371.5 | 30230488 | 9701714 | 47729573.5 | 10800579 |
| *M. midas* | Female | 13019013 | 37095320 | 12895095 | 63009428 | 23723300 |
| *M. midas* | Female | 11782515 | 23607858 | 8569692 | 43960065 | 14460645 |
| *R. metallica* | Worker | 220348.2031 | 1501856.875 | 420076.9688 | 2142282.047 | 4782087.5 |
| *R. metallica* | Worker | 250983.4688 | 1730057.875 | 329393.75 | 2310435.094 | 4289195.5 |
| *R. metallica* | Worker | 267511.9375 | 867144.9375 | 229705.7031 | 1364362.578 | 2111811.25 |
| *R. metallica* | Worker | 106794.3438 | 1468988 | 357977.6563 | 1933760 | 4333867.5 |
| *R. metallica* | Worker | 323986.1563 | 990626.4375 | 262151.3438 | 1576763.938 | 3154891.5 |
| *R. metallica* | Worker | 110373.9453 | 1139441.375 | 304612.8125 | 1554428.133 | 3688134.5 |
| *R. metallica* | Worker | 199940.3594 | 1345812.125 | 411401.2813 | 1957153.766 | 5476377 |
| *R. metallica* | Male | 934857.5 | 5907786 | 2291892.25 | 9134535.75 | 3087805.75 |
| *R. metallica* | Male | 365665.875 | 1880783.125 | 850128.625 | 3096577.625 | 983373.1875 |
| *R. metallica* | Male | 659851.375 | 4997298.5 | 1859205.625 | 7516355.5 | 3116914.25 |
| *R. metallica* | Male | 308186.5313 | 2036821.875 | 973998.0625 | 3319006.469 | 809559.8125 |
| *R. metallica* | Male | NA | 2313403.25 | 888622.5625 | NA | 1274208.375 |
| *R. metallica* | Male | 487503.375 | 5040529 | 2041047.875 | 7569080.25 | 4096356.5 |
| *R. metallica* | Female | 287359.4063 | 1999238.875 | 565267.25 | 2851865.531 | 4915800 |
| *R. metallica* | Female | 239111.75 | 1439473.5 | 426887.875 | 2105473.125 | 4148937.25 |
| *R. metallica* | Female | 181223.875 | 1631859.875 | 454519.875 | 2267603.625 | 4294643 |
| *R. metallica* | Female | 195543.75 | 1581983.5 | 455790.6875 | 2233317.938 | 4171404 |
| *R. metallica* | Female | 174171.9375 | 1411150.125 | 494490.3125 | 2079812.375 | 3919691.75 |
| *R. metallica* | Female | 227824.7969 | 1731750.625 | 509067.3438 | 2468642.766 | 5567789 |
| *R. metallica* | Female | 167689.5313 | 1722079 | 441963.9063 | 2331732.438 | 4579902 |
| *R. metallica* | Female | 313320.9063 | 1984716.5 | 528150.875 | 2826188.281 | 5027751 |

| **LIP (µm^3^)** | **COL (µm^3^)** | **PED (µm^3^)** | **MB (µm^3^)** |
| --- | --- | --- | --- |
| 23615726 | 7720040.5 | 22399396 | 53735162.5 |
| 14886048 | 6778959 | 18651800 | 40316807 |
| 19064728 | 8855096 | 20220920 | 48140744 |
| 14210554 | 7407480 | 16020762 | 37638796 |
| 62692332 | 23389496 | 48213464 | 134295292 |
| 62725600 | 24251116 | 48714868 | 135691584 |
| 72635696 | 26400286 | 48262696 | 147298678 |
| 20656056 | 5823618.5 | 12485173 | 38964847.5 |
| 56505536 | 13391906 | 27374480 | 97271922 |
| 27306970 | 7357276.5 | 17585108 | 52249354.5 |
| 9269437 | 901518.5 | 7271584.5 | 17442540 |
| 9395246 | 670143.5 | 7643628 | 17709017.5 |
| 4966651 | 489717.5313 | 3131356.75 | 8587725.281 |
| 9350075 | 608226.375 | 5159409 | 15117710.38 |
| 5845222.5 | 551693.375 | 5071441 | 11468356.88 |
| 8234714.5 | 595295.1875 | 4929773.5 | 13759783.19 |
| 11907149 | 507703.6875 | 7188683.5 | 19603536.19 |
| 2060243 | 961668.25 | 2196224.75 | 5218136 |
| 753732.125 | 291334.7813 | 720933.8125 | 1766000.719 |
| 1753287 | 769120.3125 | 2347506.5 | 4869913.813 |
| 762119.8125 | 300016.3438 | 767603.8125 | 1829739.969 |
| 1264080.875 | 564101.5 | 1146324.375 | 2974506.75 |
| 2326366.25 | 1049719.875 | 2880903.75 | 6256989.875 |
| 7174933 | 811125.1875 | 6135642 | 14121700.19 |
| 7403866.5 | 374247.6875 | 4065819 | 11843933.19 |
| 8275060.5 | 343917.7188 | 4147558.75 | 12766536.97 |
| 7156943.5 | 641365.625 | 5699993 | 13498302.13 |
| 7373506 | 573394.375 | 4995735 | 12942635.38 |
| 8593251 | 389653.3125 | 4894283.5 | 13877187.81 |
| 8523482 | 555827.375 | 5926547 | 15005856.38 |
| 10749430 | 453126.4063 | 5643912 | 16846468.41 |

| **Species** | **Caste** | **FB (µm^3^)** | **EB (µm^3^)** | **PB (µm^3^)** | **NO (µm^3^)** | **CX (µm^3^)** | **RoCB (µm^3^)** |
| --- | --- | --- | --- | --- | --- | --- | --- |
| *M. midas* | Worker | 1186463.25 | 291818.9688 | 361738.875 | 103307.8281 | 1943328.922 | 142720752 |
| *M. midas* | Worker | 1104731.875 | 421195.4375 | 440696.4688 | 109564.4766 | 2076188.258 | 142336592 |
| *M. midas* | Worker | 1341167.875 | 497124.1563 | 407898.2813 | 128954.3203 | 2375144.633 | 176023872 |
| *M. midas* | Worker | 1261718.5 | 366680.0313 | 527476.375 | 184581.5313 | 2340456.438 | 119742232 |
| *M. midas* | Worker | 1680949.75 | 640494.875 | 510406.5 | 178496.2656 | 3010347.391 | 197881856 |
| *M. midas* | Worker | 810373.1875 | 269990.7813 | 199085.7031 | 90243.92969 | 1369693.602 | 88406368 |
| *M. midas* | Worker | 1818708.375 | 591770.6875 | 541702.6875 | 189749.7188 | 3141931.469 | 142627360 |
| *M. midas* | Worker | 1660979 | 496942.9063 | 691242.5 | 236293.5625 | 3085457.969 | 161038208 |
| *M. midas* | Worker | 1636685.125 | 520027.3125 | 745071.0625 | 194913.8906 | 3096697.391 | 173082240 |
| *M. midas* | Worker | 1355154.375 | 512622.6875 | 616494.75 | 103375.4375 | 2587647.25 | 149109136 |
| *M. midas* | Worker | 1554311.625 | 425843.4688 | 461921.5938 | 126285.6172 | 2568362.305 | 162922400 |
| *M. midas* | Worker | 956714.375 | 279830.2813 | 212654.4844 | 100144.1563 | 1549343.297 | 99090968 |
| *M. midas* | Worker | 1216706.125 | 462925.875 | 678756.9375 | 171078.6719 | 2529467.609 | 135143920 |
| *M. midas* | Worker | 1250259.25 | 378103.7188 | 506782.375 | 161270.4688 | 2296415.813 | 143661040 |
| *M. midas* | Worker | 1141198.5 | 447766.25 | 493717.375 | 159612.0781 | 2242294.203 | 138721200 |
| *M. midas* | Worker | 1455409.375 | 455111.0938 | 517498.5313 | 134397.0625 | 2562416.063 | 162550080 |
| *M. midas* | Worker | 1879540 | 561579.0625 | 583881.875 | 161715.4531 | 3186716.391 | 155072192 |
| *M. midas* | Worker | 1630084.625 | 336999.1875 | 529073.75 | 142914.6875 | 2639072.25 | 165925488 |
| *M. midas* | Worker | 1452284.625 | 458034.4063 | 449664.75 | 152755.0625 | 2512738.844 | 140569696 |
| *M. midas* | Worker | 1073238.75 | 310740.375 | 414694.125 | 75001.71094 | 1873674.961 | 167904432 |
| *M. midas* | Worker | 1397851.625 | 483276.9375 | 434723.3125 | 169129.9219 | 2484981.797 | 162555248 |
| *M. midas* | Worker | 1509680.125 | 425850.1875 | 497798.6563 | 110683.6875 | 2544012.656 | 176412896 |
| *M. midas* | Worker | 1331517.375 | 373844.8438 | 417822.3438 | 130322.0156 | 2253506.578 | 145300128 |
| *M. midas* | Worker | 1327049.375 | 451071.8125 | 597461.625 | 167773.0938 | 2543355.906 | 164613472 |
| *M. midas* | Worker | 1419060.625 | 483388.9688 | 386510.1875 | 141376.9219 | 2430336.703 | 173263840 |
| *M. midas* | Worker | 1169677.375 | 325520.5 | 484684.5625 | 132671.0313 | 2112553.469 | 113241296 |
| *M. midas* | Worker | 2233649.5 | 568989.8125 | 739589.625 | 244918.2969 | 3787147.234 | 184516896 |
| *M. midas* | Worker | 1722822.375 | 548737.6875 | 717491.5 | 170315.8438 | 3159367.406 | 175452592 |
| *M. midas* | Male | 1275647.5 | 323269.8125 | 203956.8594 | 401366.2188 | 2204240.391 | 101308176 |
| *M. midas* | Male | 1215821 | 381079.8438 | 419526.75 | 201588.0938 | 2218015.688 | 105572760 |
| *M. midas* | Male | 1039779.125 | 321071.0938 | 385999.5938 | 136732.4844 | 1883582.297 | 125366024 |

| **TN (µm^3^)** | **Head**  **width (mm)** |
| --- | --- |
| 333129858.9 | 3.226 |
| 319014464.3 | 3.146 |
| 358822846.6 | 3.226 |
| 258408928.4 | 2.191 |
| 401017434.9 | 3.853 |
| 211653344.1 | 3.646 |
| 308073660.5 | 3.635 |
| 344372916.5 | 3.798 |
| 375100513.4 | 3.314 |
| 313320408.8 | 3.699 |
| 363727478.3 | 3.217 |
| 240679497.3 | 3.584 |
| 330115111.6 | 2.867 |
| 350773315.8 | 3.441 |
| 345851377.2 | 3.517 |
| 391890323.1 | 3.414 |
| 343040157.4 | 3.672 |
| 391454343.3 | 4.236 |
| 301715147.8 | 3.945 |
| 378578336 | 2.576 |
| 358314052.8 | 2.727 |
| 384866408.7 | 3.224 |
| 334153087.6 | 3.752 |
| 380750455.9 | 4.017 |
| 419336554.7 | 3.143 |
| 259798250.5 | 3.394 |
| 384056930.2 | 4.338 |
| 373609850.9 | 3.818 |
| 218418090.9 | NA |
| 228039414.7 | 2.526 |
| 274935145.3 | 2.624 |

| **Species** | **Caste** | **FB (µm^3^)** | **EB (µm^3^)** | **PB (µm^3^)** | **NO (µm^3^)** | **CX (µm^3^)** | **RoCB (µm^3^)** |
| --- | --- | --- | --- | --- | --- | --- | --- |
| *M. midas* | Male | 1761305.375 | 541609.3125 | 322419.4688 | 255243.6875 | 2880577.844 | 127187720 |
| *M. midas* | Male | 1508976 | 461746.5938 | 432899 | 172003.3281 | 2575624.922 | 125793104 |
| *M. midas* | Male | 1282037.625 | 349495.9375 | 447385.375 | 127960.4219 | 2206879.359 | 124330928 |
| *M. midas* | Male | 1356639.125 | 308198.2813 | 351412.125 | 126949.4375 | 2143198.969 | 110900928 |
| *M. midas* | Female | 1470281.125 | 475740.0313 | 469180.375 | 168035.4844 | 2583237.016 | 218436368 |
| *M. midas* | Female | 1608010.5 | 620569.8125 | 638932.1875 | 195097.0313 | 3062609.531 | 190741648 |
| *M. midas* | Female | 1958317.625 | 590075.625 | 625142.8125 | 183341.3594 | 3356877.422 | 267184304 |
| *M. midas* | Female | 759876 | 194287.4375 | 251453.6719 | 59401.375 | 1265018.484 | 90746400 |
| *M. midas* | Female | 1544096.75 | 342682.9063 | 746875.0625 | 225850.3281 | 2859505.047 | 138356272 |
| *M. midas* | Female | 1007893.938 | 202675.2031 | 377454.5938 | 132864.3281 | 1720888.063 | 85156880 |
| *R. metallica* | Worker | 306011.75 | 52865.23047 | 92810.98438 | 35408.06641 | 487096.0313 | 22896840 |
| *R. metallica* | Worker | 212295.5625 | 57414.67188 | 81008.28906 | 35243.48438 | 385962.0078 | 23309040 |
| *R. metallica* | Worker | 53582.32422 | 16410.91016 | 53887.97266 | 13848.17383 | 137729.3809 | 10154191 |
| *R. metallica* | Worker | 155559.7813 | 69499.49219 | 76876.17969 | 20831.03906 | 322766.4922 | 18195572 |
| *R. metallica* | Worker | 107329.2266 | 40016.28516 | 61435.10938 | 19091.20117 | 227871.8223 | 13435997 |
| *R. metallica* | Worker | 232021.5781 | 51398.71094 | 91855.83594 | 24298.96094 | 399575.0859 | 17661196 |
| *R. metallica* | Worker | 266736.0625 | 83853.16406 | 116110.7109 | 26979.25391 | 493679.1914 | 25501214 |
| *R. metallica* | Male | 333629.6875 | 77414.97656 | 135472.2813 | 67473.60156 | 613990.5469 | 16586475 |
| *R. metallica* | Male | 109747.9531 | 30929.15625 | 37447.67188 | 19546.73242 | 197671.5137 | 6322837.5 |
| *R. metallica* | Male | 259494.5625 | 71665.47656 | 80696.76563 | 35549.13281 | 447405.9375 | 15462521 |
| *R. metallica* | Male | 113797.7813 | 28639.74023 | 34908.44922 | 13607.18359 | 190953.1543 | 5921270 |
| *R. metallica* | Male | 126026.6172 | 38123.625 | 46305.56641 | 25001.36133 | 235457.1699 | 7659186.5 |
| *R. metallica* | Male | 380236.625 | 108578.6797 | 148633.2344 | 69739.69531 | 707188.2344 | 17628956 |
| *R. metallica* | Female | 244607.9531 | 50831.49609 | 153920.8438 | 34573.41016 | 483933.7031 | 23568494 |
| *R. metallica* | Female | 179585.8281 | 33502.32813 | 95199.98438 | 20931.54297 | 329219.6836 | 18147070 |
| *R. metallica* | Female | 192371.5625 | 47844.58594 | 70073.23438 | NA | NA | 19307082 |
| *R. metallica* | Female | 281219.0313 | 55228.11719 | 114794.0781 | 46305.56641 | 497546.793 | 20068990 |
| *R. metallica* | Female | 247539.0469 | 56062.76953 | 129500.4141 | 39028.80859 | 472131.0391 | 19561310 |
| *R. metallica* | Female | 195684.8125 | 52559.58203 | 128030.9531 | 31364.11719 | 407639.4648 | 21557116 |
| *R. metallica* | Female | 275786.4375 | 67019.48438 | 150071.1563 | 46495.60156 | 539372.6797 | 22433026 |
| *R. metallica* | Female | 286674.6875 | 66485.82031 | 181127.5313 | 50097.83984 | 584385.8789 | 21609550 |

| **TN (µm^3^)** | **Head**  **width (mm)** |
| --- | --- |
| 279617145.3 | NA |
| NA | 2.711 |
| 258765580.4 | 2.689 |
| 232026371 | 2.514 |
| 477754765 | 4.531 |
| 455409039.5 | 4.59 |
| 553693483.4 | 4.666 |
| 189506418.5 | NA |
| 325220427 | NA |
| 197547832.6 | NA |
| 47750845.58 | 1.65 |
| 48003650.1 | 1.683 |
| 22355819.49 | 1.376 |
| 39903676.37 | NA |
| 29863881.13 | 1.481 |
| 37063116.91 | 1.436 |
| 53031960.14 | 1.532 |
| 34640943.05 | 1.281 |
| 12366460.54 | 1.158 |
| 31413110.5 | 1.289 |
| 12070529.4 | 1.293 |
| NA | 1.189 |
| 36258570.86 | NA |
| 45941793.42 | 1.672 |
| 36574633.25 | 1.592 |
| NA | 1.576 |
| 40469560.86 | 1.714 |
| 38975580.54 | 1.684 |
| 43878375.04 | 1.521 |
| 44889889.49 | NA |
| 46894343.57 | NA |

LA = lamina; ME = medulla; LO = lobula; OL = optic lobe; AL = antennal lobe; LIP = calyx lip; COL = calyx collar; PED = peduncle and lobes; MB = mushroom body; FB = fan-shaped body;

EB =ellipsoid body; PB = protocerebral bridge; NO = noduli; CX = central complex; RoCB = rest of the central brain; TN = total neuropil; HW = head width
