## Supplementary material for "Caste- and Sex-Specific Differential Investment in Brain Regions of Australian Ants": Table S2

| **Caste** | **y** | **Do groups have a common slope? H0 = slopes are equal** | **Is the common slope different from 1 H0 = common slope not different from one** | **Is the linear model (lm) for *M. midas* significant?** | **Is the linear model (lm) for *R. metallica* significant?** | **Are there differences in elevation? H0 : no difference in elevation** | **α*_M.midas_*** | **α*_R. metallica_*** | **gsi  (*M. midas* 'M'/ *R. metallica* 'R')** |
| --- | --- | --- | --- | --- | --- | --- | --- | --- | --- |
| Workers | LA | Yes p = 0.73 | No p = 0.08 | No p = 0.21 | No p = 0.53 | No p = 0.21 | -5.17 | -5.52 | 2.21 |
|  | ME | Yes p = 0.15 | No p = 0.37 | Yes p = 0.01 | Yes p = 0.02 | Yes p = 4.42 x 10^-06^ | -0.24 | -0.8 | 3.62 |
|  | LO | No p = 0.05 | No p = 0.15 | Yes p = 0.01 | Yes p = 0.01 | Yes p = 6.78 x 10^-12^ | -0.2 | -0.96 | 5.76 |
|  | OL | No p = 0.03 |  | Yes p = 0.02 | Yes p = 0.01 |  |  |  |  |
|  | AL | Yes p = 0.26 | No p = 0.41 | Yes p < 0.01 | Yes p < 0.01 | Yes p =0.41 x 10^-3^ | -1.3 | -1.03 | 0.53 |
|  | LIP | Yes p = 0.74 | No p = 0.79 | Yes p < 0.01 | Yes p < 0.01 | No p = 0.33 | 0.009 | 0.09 | 0.84 |
|  | COL | Yes p = 0.45 | No p = 0.26 | Yes p < 0.01 | No p = 0.30 | Yes p = 7.50 x 10^-12^ | 0.48 | -0.22 | 4.93 |
|  | PED | Yes p = 0.85 | No p = 0.61 | Yes p < 0.01 | Yes p < 0.01 | No p = 0.97 | 0.002 | -4E-04 | 1.01 |
|  | MB | Yes p = 0.94 | Yes p = 0.03 | Yes p < 0.01 | Yes p < 0.01 | Yes p = 0.02 | 0.92 | 0.85 | 1.19 |
|  | FB | No p = 0.05 | Yes p = 0.02 | Yes p < 0.01 | Yes p = 0.01 | Yes p = 0.01 | -5.4 | -5.02 | 0.41 |
|  | EB | Yes p = 0.39 | No p = 0.05 | Yes p < 0.01 | Yes p = 0.02 | Yes p = 0.04 | -5.71 | -5.38 | 0.47 |
|  | PB | No p = 0.001 |  | Yes p < 0.01 | Yes p = 0.01 |  |  |  |  |
|  | NO | Yes p = 0.14 | No p = 0.08 | No p = 0.07 | Yes p = 0.01 | Yes p = 0.02 | -4.99 | -4.62 | 0.43 |
|  | CX | Yes p = 0.43 | No p = 0.06 | Yes p < 0.01 | Yes p < 0.01 | Yes p = 0.005 | -4.27 | -3.95 | 0.48 |

Supplementary Table 2: Outputs of standardized major axis regression (SMA) on log transformed data (log y = α + βlog x) comparing the scaling relationships of each neuropil with the volume of the reference structure, the RoCB, between the two species, *Myrmecia midas* and *Rhytidoponera metallica*. Abbreviations and gsi conventions as described in ‘Methods’ section

Abbreviations and gsi conventions as described in 'Methods' section

| **Caste** | **y** | **Do groups have a common slope? H0 = slopes are equal** | **Is the common slope different from 1 H0 = common slope not different from one** | **Is the lm for M. midas significant?** | **Is the lm for R. metallica significant?** | **Are there differences in elevation? H0 : no difference in elevation** | **α*_M.midas_*** | **α*_R. metallica_*** | **gsi  (*M. midas* 'M'/ *R. metallica* 'R')** |
| --- | --- | --- | --- | --- | --- | --- | --- | --- | --- |
| Alate females | LA | No p = 0.02 |  | Yes p = 0.01 | No p = 0.34 |  |  |  |  |
|  | ME | Yes p = 0.05 | No p = 0.13 | Yes p < 0.01 | Yes p = 0.01 | Yes p = 4.5x 10^-14^ | 0.1 | -0.5 | 4.05 |
|  | LO | Yes p = 0.36 | No p = 0.22 | Yes p < 0.01 | No p = 0.06 | Yes p = < 2.22x 10^-16^ | 0.28 | -0.46 | 5.57 |
|  | OL | Yes  p = 0.07 | No p = 0.09 | Yes p < 0.01 | Yes p = 0.02 | Yes p = < 2.22x 10^-16^ | 0.9 | 0.16 | 5.53 |
|  | AL | Yes p = 0.31 | No p = 0.57 | Yes p = 0.04 | No p = 0.07 | No p = 0.18 | -1.25 | -1 | 0.57 |
|  | LIP | Yes p = 0.43 | No p = 0.49 | Yes p = 0.01 | No p = 0.55 | No p = 0.15 | -1.94 | -1.68 | 0.55 |
|  | COL | No p = 0.03 |  | Yes p < 0.01 | No p = 0.12 |  |  |  |  |
|  | PED | Yes p = 0.22 | No p = 0.05 | Yes p = 0.01 | Yes p = 0.01 | Yes p = 0.01 | -4 | -3.55 | 0.36 |
|  | MB | Yes p = 0.91 | No p = 0.48 | Yes p < 0.01 | No p = 0.06 | No p = 0.13 | -1.86 | -1.64 | 0.6 |
|  | FB | No  p = 0.02 |  | Yes p = 0.02 | No p = 0.22 |  |  |  |  |
|  | EB | No  p = 0.04 |  | Yes p = 0.01 | No p = 0.11 |  |  |  |  |
|  | PB | No  p = 0.01 |  | No p = 0.27 | Yes p = 0.03 |  |  |  |  |
|  | NO | Yes  p = 0.05 | Yes  p = 0.04 | No p = 0.26 | No p = 0.36 | Yes p = 0.04 | -10.18 | -9.17 | 0.1 |
|  | CX | Yes p = 0.05 | No p = 0.13 | Yes p = 0.03 | No p = 0.17 | No p = 0.29 | -2.55 | -2.31 | 0.58 |
| **Caste** | **y** | **Do groups have a common slope? H0 = slopes are equal** | **Is the common slope different from 1 H0 = common slope not different from one** | **Is the lm for M. midas significant?** | **Is the lm for R. metallica significant?** | **Are there differences in elevation? H0 : no difference in elevation** | **α*_M.midas_*** | **α*_R. metallica_*** | **gsi  (*M. midas* 'M'/ *R. metallica* 'R')** |
| Alate males | LA | Yes p = 0.36 | No p = 0.65 | No p = 0.13 | No p = 0.16 | No p = 0.13 | -1.02 | -1.43 | 2.54 |
|  | ME | Yes p = 0.1 | No p = 0.22 | Yes p = 0.02 | Yes p < 0.01 | No p = 0.81 | -0.93 | -0.9 | 0.95 |
|  | LO | Yes p = 0.13 | No p = 0.30 | No p = 0.17 | Yes p < 0.01 | No p = 0.87 | -0.49 | -0.51 | 1.05 |
|  | OL | Yes  p = 0.12 | No p = 0.29 | Yes p = 0.03 | Yes p = 0.01 | No p = 0.44 | -0.12 | -0.2 | 1.22 |
|  | AL | Yes  p = 0.89 | Yes  p = 0.02 | No p = 0.95 | Yes p < 0.01 | Yes p = 2.26 x 10^-07^ | -3.65 | -3.21 | 0.36 |
|  | LIP | Yes  p = 0.19 | No p = 0.43 | No p = 0.89 | Yes p < 0.01 | No p = 0.59 | -0.84 | -0.92 | 1.19 |
|  | COL | No p = 0.03 |  | Yes p = 0.03 | Yes p < 0.01 |  |  |  |  |
|  | PED | Yes  p = 0.42 | No p = 0.13 | No p = 0.05 | Yes p < 0.01 | No p = 0.11 | -2.43 | -2.26 | 0.68 |
|  | MB | Yes  p = 0.56 | No p = 0.6 | No p = 0.17 | Yes p < 0.01 | No p = 0.72 | -1.17 | -1.13 | 0.91 |
|  | FB | Yes  p = 0.34 | No p = 0.27 | No p = 0.45 | Yes p < 0.01 | Yes p = 0.001 | -2.99 | -2.65 | 0.45 |
|  | EB | Yes  p = 0.11 | No p = 0.15 | No p = 0.23 | Yes p < 0.01 | Yes p = 0.006 | -3.37 | -3.07 | 0.5 |
|  | PB | Yes  p = 0.07 | Yes p = 0.02 | No p = 0.22 | Yes p = 0.01 | Yes p = 1.68 x 10^-4^ | -5.22 | -4.54 | 0.21 |
|  | NO | No p = 0.01 |  | No p = 0.22 | Yes p = 0.01 |  |  |  |  |
|  | CX | Yes p = 0.63 | No p = 0.33 | No p = 0.44 | Yes p < 0.01 | Yes p = 4.68 x 10^-4^ | -2.96 | -2.57 | 0.41 |
