## Supplementary material for "Caste- and Sex-Specific Differential Investment in Brain Regions of Australian Ants": Table S3

| **Species** | **Region of interest (y)**  Supplementary Table 3: Outputs of standardized major axis regression (SMA) on log transformed data (log y = α + βlog x) comparing the scaling relationships of each neuropil with the volume of the reference structure, the RoCB, between the workers, alate males (males) and alate females (females) of *Myrmecia midas* and *Rhytidoponera metallica*. Abbreviations and gsi conventions as described in ‘Methods’ section | | **Do groups have a common slope? H0 = slopes are equal** | | | **Pair wise difference in slope Does slope of caste 1 differ from slope of caste 2? H0 = no difference in elevation** | | | | | | | | **Is the common slope different from 1? H0 = common slope not different from 1** | | | **Is the linear model significant? (p-values) Yes, p < 0.05 No, p > 0.05** | | | | **Are there differences in elevation? H0 : no difference in elevation** | | | | **Pair wise difference in elevation Does elevation of caste 1 differ from elevation of caste 2? H0 = no difference in elevation** | | | | | | | | | **α_males_** | | | **α_Workers_** | | | **α_Females_** | | | **gsi (m/w)** | | | **gsi (f/m)** | | | **gsi (w/f)** |
| --- | --- | --- | --- | --- | --- | --- | --- | --- | --- | --- | --- | --- | --- | --- | --- | --- | --- | --- | --- | --- | --- | --- | --- | --- | --- | --- | --- | --- | --- | --- | --- | --- | --- | --- | --- | --- | --- | --- | --- | --- | --- | --- | --- | --- | --- | --- | --- | --- | --- |
|  |  |  |  |  |  | *Caste 1* | | *Caste 2* | | | *p-value* | | |  | | |  | | | |  | | | | *Caste 1* | | | *Caste 2* | | | *p-value* | | |  | | |  | | |  | | |  | | |  | | |  |
| *M. midas* | LA | | Yes p = 0.16 | | |  | |  | | |  | | | No p = 0.11 | | | Males = 0.13 Workers = 0.21 Females = 0.01 | | | | Yes p = 2.90 x 10^-11^ | | | | Males | | | Workers | | | 8.71 x 10^-11^ | | | -2.72 | | | -2.99 | | | -2.84 | | | 1.84 | | | 0.77 | | | 0.71 |
|  |  |  |  |  |  |  | |  | | |  | | |  |  |  |  |  |  |  |  |  |  |  | Females | | | Males | | | 0.07 | | |  |  |  |  |  |  |  |  |  |  |  |  |  |  |  |  |
|  |  |  |  |  |  |  | |  | | |  | | |  |  |  |  |  |  |  |  |  |  |  | Workers | | | Females | | | 0.009 | | |  |  |  |  |  |  |  |  |  |  |  |  |  |  |  |  |
|  | ME | | Yes  p = 0.08 | | |  | |  | | |  | | | No p = 0.18 | | | Males = 0.02 Workers = 0.01 Females < 0.01 | | | | Yes p = 8.37 x 10^-12^ | | | | Males | | | Workers | | | 2.03 x 10^-12^ | | | -0.24 | | | -0.42 | | | -0.33 | | | 1.5 | | | 0.81 | | | 0.82 |
|  |  |  |  |  |  |  | |  | | |  | | |  |  |  |  |  |  |  |  |  |  |  | Females | | | Males | | | 0.002 | | |  |  |  |  |  |  |  |  |  |  |  |  |  |  |  |  |
|  |  |  |  |  |  |  | |  | | |  | | |  |  |  |  |  |  |  |  |  |  |  | Workers | | | Females | | | 1.87 x 10^-4^ | | |  |  |  |  |  |  |  |  |  |  |  |  |  |  |  |  |
|  | LO | | Yes  p = 0.08 | | |  | |  | | |  | | | No p = 0.16 | | | Males = 0.17 Workers = 0.01 Females < 0.01 | | | | Yes p = 9.68 x 10^-7^ | | | | Males | | | Workers | | | 1.53 x 10^-7^ | | | -0.09 | | | -0.24 | | | -0.18 | | | 1.42 | | | 0.81 | | | 0.87 |
|  |  |  |  |  |  |  | |  | | |  | | |  |  |  |  |  |  |  |  |  |  |  | Females | | | Males | | | 0.002 | | |  |  |  |  |  |  |  |  |  |  |  |  |  |  |  |  |
|  |  |  |  |  |  |  | |  | | |  | | |  |  |  |  |  |  |  |  |  |  |  | Workers | | | Females | | | 0.005 | | |  |  |  |  |  |  |  |  |  |  |  |  |  |  |  |  |
|  | OL | | Yes p = 0.06 | | |  | |  | | |  | | | No p = 0.12 | | | Males = 0.03 Workers = 0.02 Females <0.01 | | | | Yes p =1.01 x 10^-12^ | | | | Males | | | Workers | | | 8.68 x 10^-13^ | | | 0.66 | | | 0.49 | | | 0.58 | | | 1.51 | | | 0.82 | | | 0.81 |
|  |  |  |  |  |  |  | |  | | |  | | |  |  |  |  |  |  |  |  |  |  |  | Females | | | Males | | | 3.48 x 10^-4^ | | |  |  |  |  |  |  |  |  |  |  |  |  |  |  |  |  |
|  |  |  |  |  |  |  | |  | | |  | | |  |  |  |  |  |  |  |  |  |  |  | Workers | | | Females | | | 3.56 x 10^-6^ | | |  |  |  |  |  |  |  |  |  |  |  |  |  |  |  |  |
| *R. metallica* | LA | | Yes  p = 0.12 | | |  | |  | | |  | | | No p = 0.13 | | | Males = 0.16 Workers = 0.53 Females = 0.34 | | | | Yes p = 3.55 x 10^-7^ | | | | Males | | | Workers | | | 7.12 x 10^-6^ | | | -3.73 | | | -4.42 | | | -4.46 | | | 4.91 | | | 0.19 | | | 1.09 |
|  |  |  |  |  |  |  | |  | | |  | | |  |  |  |  |  |  |  |  |  |  |  | Females | | | Males | | | 7.89 x 10^-7^ | | |  |  |  |  |  |  |  |  |  |  |  |  |  |  |  |  |
|  |  |  |  |  |  |  | |  | | |  | | |  |  |  |  |  |  |  |  |  |  |  | Workers | | | Females | | | 0.64 | | |  |  |  |  |  |  |  |  |  |  |  |  |  |  |  |  |
|  | ME | | Yes  p = 0.13 | | |  | |  | | |  | | | No p = 0.25 | | | Males < 0.01 Workers = 0.02 Females = 0.01 | | | | Yes p < 2.22 x 10^-16^ | | | | Males | | | Workers | | | < 2.22 x 10^-16^ | | | -0.65 | | | -1.31 | | | -1.25 | | | 4.57 | | | 0.25 | | | 0.87 |
|  |  |  |  |  |  |  | |  | | |  | | |  |  |  |  |  |  |  |  |  |  |  | Females | | | Males | | | < 2.22 x 10^-16^ | | |  |  |  |  |  |  |  |  |  |  |  |  |  |  |  |  |
|  |  |  |  |  |  |  | |  | | |  | | |  |  |  |  |  |  |  |  |  |  |  | Workers | | | Females | | | 0.09 | | |  |  |  |  |  |  |  |  |  |  |  |  |  |  |  |  |
|  | LO | | Yes  p = 0.33 | | |  | |  | | |  | | | No p = 0.26 | | | Males < 0.01 Workers = 0.01 Females = 0.06 | | | | Yes p < 2.22 x 10^-16^ | | | | Males | | | Workers | | | < 2.22 x 10^-16^ | | | 0.17 | | | -0.66 | | | -0.54 | | | 6.66 | | | 0.2 | | | 0.76 |
|  |  |  |  |  |  |  | |  | | |  | | |  |  |  |  |  |  |  |  |  |  |  | Females | | | Males | | | < 2.22 x 10^-16^ | | |  |  |  |  |  |  |  |  |  |  |  |  |  |  |  |  |
|  |  |  |  |  |  |  | |  | | |  | | |  |  |  |  |  |  |  |  |  |  |  | Workers | | | Females | | | 2.26 x 10^-8^ | | |  |  |  |  |  |  |  |  |  |  |  |  |  |  |  |  |
|  | OL | | Yes  p = 0.06 | | |  | |  | | |  | | | No p = 0.11 | | | Males = 0.01 Workers = 0.01 Females = 0.02 | | | | Yes p < 2.22 x 10^-16^ | | | | Males | | | Workers | | | < 2.22 x 10^-16^ | | | 0.24 | | | -0.44 | | | -0.38 | | | 4.8 | | | 0.24 | | | 0.87 |
|  |  |  |  |  |  |  | |  | | |  | | |  |  |  |  |  |  |  |  |  |  |  | Females | | | Males | | | < 2.22 x 10^-16^ | | |  |  |  |  |  |  |  |  |  |  |  |  |  |  |  |  |
|  |  |  |  |  |  |  | |  | | |  | | |  |  |  |  |  |  |  |  |  |  |  | Workers | | | Females | | | 0.02 | | |  |  |  |  |  |  |  |  |  |  |  |  |  |  |  |  |
| **Species** | **ROI (y)** | | | **Common slope?** | | | **Pair wise difference in slope** | | | | | | | | | **Common slope different from 1?** | | | **Is the linear model significant?** | | | **Are there differences in elevation?** | | **Pair wise difference in elevation** | | | | | | | | | **α_males_** | | | **α_Workers_** | | | **α_Females_** | | | **gsi (m/w)** | | | **gsi (f/m)** | | | **gsi (w/f)** | |
| *M. midas* | AL | | | Yes p = 0.6 | | |  | | |  | | |  | | | No p = 0.49 | | | Males = 0.95 Workers < 0.01 Females = 0.04 | | | Yes p = 0.04 | | Males | | | Workers | | | 0.01 | | | -1.71 | | | -1.79 | | | -1.75 | | | 1.22 | | | 0.9 | | | 0.91 | |
|  |  |  |  |  |  |  |  | | |  | | |  | | |  |  |  |  |  |  |  |  | Females | | | Males | | | 0.56 | | |  |  |  |  |  |  |  |  |  |  |  |  |  |  |  |  |  |
|  |  |  |  |  |  |  |  | | |  | | |  | | |  |  |  |  |  |  |  |  | Workers | | | Females | | | 0.43 | | |  |  |  |  |  |  |  |  |  |  |  |  |  |  |  |  |  |
| *R. metallica* | AL | | | Yes  p = 0.11 | | |  | | |  | | |  | | | Yes p = 0.03 | | | Males < 0.01 Workers < 0.01 Females = 0.07 | | | No p = 0.97 | |  | | |  | | |  | | | -2.94 | | | -2.94 | | | -2.94 | | | 0.98 | | | 1 | | | 1.02 | |
| *M. midas* | LIP | | | Yes  p = 0.4 | | |  | | |  | | |  | | | No p = 0.59 | | | Males = 0.89 Workers < 0.01 Females = 0.01 | | | Yes p = 9.99 x 10^-16^ | | Males | | | Workers | | | < 2.22 x 10^-16^ | | | -1.25 | | | -0.92 | | | -0.97 | | | 0.47 | | | 1.92 | | | 1.11 | |
|  |  |  |  |  |  |  |  | | |  | | |  | | |  |  |  |  |  |  |  |  | Females | | | Males | | | 1.13 x 10^-5^ | | |  |  |  |  |  |  |  |  |  |  |  |  |  |  |  |  |  |
|  |  |  |  |  |  |  |  | | |  | | |  | | |  |  |  |  |  |  |  |  | Workers | | | Females | | | 0.27 | | |  |  |  |  |  |  |  |  |  |  |  |  |  |  |  |  |  |
|  | COL | | | No  p = 0.002 | | | Males | | | Workers | | | 0.005 | | |  | | | Males = 0.03 Workers < 0.01 Females < 0.01 | | |  | |  | | |  | | |  | | |  | | |  | | |  | | |  | | |  | | |  | |
|  |  |  |  |  |  |  | Females | | | Males | | | 0.08 | | |  | | |  |  |  |  | |  | | |  | | |  | | |  | | |  | | |  | | |  | | |  | | |  | |
|  |  |  |  |  |  |  | Workers | | | Females | | | 0.02 | | |  | | |  |  |  |  | |  | | |  | | |  | | |  | | |  | | |  | | |  | | |  | | |  | |
|  | PED | | | Yes  p = 0.13 | | |  | | |  | | |  | | | No p = 0.26 | | | Males = 0.05 Workers < 0.01 Females = 0.01 | | | Yes p < 2.22 x 10^-16^ | | Males | | | Workers | | | < 2.22 x 10^-16^ | | | -0.98 | | | -0.74 | | | -0.88 | | | 0.58 | | | 1.27 | | | 1.37 | |
|  |  |  |  |  |  |  |  | | |  | | |  | | |  |  |  |  |  |  |  |  | Females | | | Males | | | 0.13 | | |  |  |  |  |  |  |  |  |  |  |  |  |  |  |  |  |  |
|  |  |  |  |  |  |  |  | | |  | | |  | | |  |  |  |  |  |  |  |  | Workers | | | Females | | | 0.002 | | |  |  |  |  |  |  |  |  |  |  |  |  |  |  |  |  |  |
|  | MB | | | Yes   p = 0.21 | | |  | | |  | | |  | | | No  p = 0.33 | | | Males = 0.17 Workers < 0.01 Females < 0.01 | | | Yes p < 2.22 x 10^-16^ | | Males | | | Workers | | | < 2.22 x 10^-16^ | | | -0.22 | | | 0.07 | | | -0.01 | | | 0.51 | | | 1.63 | | | 1.21 | |
|  |  |  |  |  |  |  |  | | |  | | |  | | |  |  |  |  |  |  |  |  | Females | | | Males | | | 5.07 x 10^-5^ | | |  |  |  |  |  |  |  |  |  |  |  |  |  |  |  |  |  |
|  |  |  |  |  |  |  |  | | |  | | |  | | |  |  |  |  |  |  |  |  | Workers | | | Females | | | 0.04 | | |  |  |  |  |  |  |  |  |  |  |  |  |  |  |  |  |  |
| *R. metallica* | LIP | | | Yes  p = 0.46 | | |  | | |  | | |  | | | No p = 0.64 | | | Males < 0.01 Workers < 0.01 Females = 0.55 | | | Yes p < 2.22 x 10^-16^ | | Males | | | Workers | | | < 2 x 10^-16^ | | | -0.61 | | | -0.06 | | | -0.12 | | | 0.28 | | | 3.1 | | | 1.15 | |
|  |  |  |  |  |  |  |  | | |  | | |  | | |  |  |  |  |  |  |  |  | Females | | | Males | | | < 2 x 10^-16^ | | |  |  |  |  |  |  |  |  |  |  |  |  |  |  |  |  |  |
|  |  |  |  |  |  |  |  | | |  | | |  | | |  |  |  |  |  |  |  |  | Workers | | | Females | | | 0.03 | | |  |  |  |  |  |  |  |  |  |  |  |  |  |  |  |  |  |
|  | COL | | | No  p = 0.01 | | | Males | | | Workers | | | 0.2 | | |  | | | Males < 0.01 Workers = 0.03 Females = 0.12 | | |  | |  | | |  | | |  | | |  | | |  | | |  | | |  | | |  | | |  | |
|  |  |  |  |  |  |  | Females | | | Males | | | 0.009 | | |  | | |  |  |  |  |  |  | | |  | | |  | | |  | | |  | | |  | | |  | | |  | | |  | |
|  |  |  |  |  |  |  | Workers | | | Females | | | 0.004 | | |  | | |  |  |  |  |  |  | | |  | | |  | | |  | | |  | | |  | | |  | | |  | | |  | |
|  | PED | | | Yes  p = 0.11 | | |  | | |  | | |  | | | No p = 0.07 | | | Males < 0.01 Workers < 0.01 Females = 0.01 | | | Yes p = 2.22 x 10^-15^ | | Males | | | Workers | | | < 2.22 x 10^-16^ | | | -2.04 | | | -1.73 | | | -1.84 | | | 0.49 | | | 1.59 | | | 1.28 | |
|  |  |  |  |  |  |  |  | | |  | | |  | | |  |  |  |  |  |  |  |  | Females | | | Males | | | 2.81 x 10^-6^ | | |  |  |  |  |  |  |  |  |  |  |  |  |  |  |  |  |  |
|  |  |  |  |  |  |  |  | | |  | | |  | | |  |  |  |  |  |  |  |  | Workers | | | Females | | | 3.56 x 10^-4^ | | |  |  |  |  |  |  |  |  |  |  |  |  |  |  |  |  |  |
|  | MB | | | Yes  p = 0.16 | | |  | | |  | | |  | | | No p = 0.09 | | | Males < 0.01 Workers < 0.01 Females = 0.06 | | | Yes p < 2.22 x 10^-16^ | | Males | | | Workers | | | < 2.22 x 10^-16^ | | | 0.28 | | | 0.7 | | | 0.62 | | | 0.38 | | | 2.21 | | | 1.18 | |
|  |  |  |  |  |  |  |  | | |  | | |  | | |  |  |  |  |  |  |  |  | Females | | | Males | | | 2.35 x 10^-12^ | | |  |  |  |  |  |  |  |  |  |  |  |  |  |  |  |  |  |
|  |  |  |  |  |  |  |  | | |  | | |  | | |  |  |  |  |  |  |  |  | Workers | | | Females | | | 6.48 x 10^-8^ | | |  |  |  |  |  |  |  |  |  |  |  |  |  |  |  |  |  |
| **Species** | **ROI (y)** | **Common slope?** | | | **Pair wise difference in slope** | | | | | | | | | | **Common slope different from 1?** | | |  | | **Are there differences in elevation?** | | | **Pair wise difference in elevation** | | | | | | | | | **α_males_** | | | **α_Workers_** | | | **α_Females_** | | | **gsi (m/w)** | | | **gsi (f/m)** | | | **gsi (w/f)** | | |
| *M. midas* | FB | Yes   p = 0.19 | | |  | | | |  | | |  | | | No  p = 0.24 | | | Males = 0.45 Workers < 0.01 Females = 0.02 | | Yes p = 0.01 | | | Males | | | Workers | | | 0.002 | | | -2.8 | | | -2.9 | | | -2.92 | | | 1.27 | | | 0.75 | | | 1.05 | | |
|  |  |  |  |  |  | | | |  | | |  | | |  |  |  |  |  |  |  |  | Females | | | Males | | | 0.05 | | |  |  |  |  |  |  |  |  |  |  |  |  |  |  |  |  |  |  |
|  |  |  |  |  |  | | | |  | | |  | | |  |  |  |  |  |  |  |  | Workers | | | Females | | | 0.67 | | |  |  |  |  |  |  |  |  |  |  |  |  |  |  |  |  |  |  |
|  | EB | Yes   p = 0.3 | | |  | | | |  | | |  | | | No  p = 0.1 | | | Males = 0.24 Workers < 0.01 Females = 0.01 | | Yes p = 0.01 | | | Males | | | Workers | | | 0.03 | | | -4.31 | | | -4.39 | | | -4.46 | | | 1.18 | | | 0.71 | | | 1.2 | | |
|  |  |  |  |  |  | | | |  | | |  | | |  |  |  |  |  |  |  |  | Females | | | Males | | | 0.004 | | |  |  |  |  |  |  |  |  |  |  |  |  |  |  |  |  |  |  |
|  |  |  |  |  |  | | | |  | | |  | | |  |  |  |  |  |  |  |  | Workers | | | Females | | | 0.03 | | |  |  |  |  |  |  |  |  |  |  |  |  |  |  |  |  |  |  |
|  | PB | Yes   p = 0.14 | | |  | | | |  | | |  | | | Yes  p = 0.001 | | | Males = 0.22 Workers < 0.01 Females = 0.27 | | No p = 0.64 | | | Males | | | Workers | | | 0.74 | | | -8.07 | | | -8.12 | | | -8.12 | | | 1.12 | | | 0.88 | | | 1.01 | | |
|  |  |  |  |  |  | | | |  | | |  | | |  |  |  |  |  |  |  |  | Females | | | Males | | | 0.22 | | |  |  |  |  |  |  |  |  |  |  |  |  |  |  |  |  |  |  |
|  |  |  |  |  |  | | | |  | | |  | | |  |  |  |  |  |  |  |  | Workers | | | Females | | | 0.97 | | |  |  |  |  |  |  |  |  |  |  |  |  |  |  |  |  |  |  |
|  | NO | Yes p = 0.05 | | |  | | | |  | | |  | | | Yes  p = 0.003 | | | Males = 0.22 Workers = 0.07 Females = 0.26 | | Yes p = 0.02 | | | Males | | | Workers | | | 0.003 | | | -7.93 | | | -8.22 | | | -8.21 | | | 1.96 | | | 0.52 | | | 0.98 | | |
|  |  |  |  |  |  | | | |  | | |  | | |  |  |  |  |  |  |  |  | Females | | | Males | | | 0.09 | | |  |  |  |  |  |  |  |  |  |  |  |  |  |  |  |  |  |  |
|  |  |  |  |  |  | | | |  | | |  | | |  |  |  |  |  |  |  |  | Workers | | | Females | | | 0.93 | | |  |  |  |  |  |  |  |  |  |  |  |  |  |  |  |  |  |  |
|  | CX | Yes  p = 0.4 | | |  | | | |  | | |  | | | No  p = 0.4 | | | Males = 0.44 Workers < 0.01 Females = 0.03 | | Yes p = 0.02 | | | Males | | | Workers | | | 0.004 | | | -2.68 | | | -2.77 | | | -2.8 | | | 1.23 | | | 0.77 | | | 1.06 | | |
|  |  |  |  |  |  | | | |  | | |  | | |  |  |  |  |  |  |  |  | Females | | | Males | | | 0.09 | | |  |  |  |  |  |  |  |  |  |  |  |  |  |  |  |  |  |  |
|  |  |  |  |  |  | | | |  | | |  | | |  |  |  |  |  |  |  |  | Workers | | | Females | | | 0.65 | | |  |  |  |  |  |  |  |  |  |  |  |  |  |  |  |  |  |  |
| *R. metallica* | FB | No  p = 0.03 | | | Males | | | | Workers | | | 0.02 | | |  | | | Males < 0.01 Workers = 0.01 Females = 0.22 | |  | | |  | | |  | | |  | | |  | | |  | | |  | | |  | | |  | | |  | | |
|  |  |  |  |  | Females | | | | Males | | | 0.09 | | |  | | |  |  |  |  |  |  | | |  | | |  | | |  | | |  | | |  | | |  | | |  | | |  | | |
|  |  |  |  |  | Workers | | | | Females | | | 0.69 | | |  | | |  |  |  |  |  |  | | |  | | |  | | |  | | |  | | |  | | |  | | |  | | |  | | |
|  | EB | No  p = 0.04 | | | Males | | | | Workers | | | 0.13 | | |  | | | Males < 0.01 Workers = 0.02 Females = 0.11 | |  | | |  | | |  | | |  | | |  | | |  | | |  | | |  | | |  | | |  | | |
|  |  |  |  |  | Females | | | | Males | | | 0.03 | | |  | | |  |  |  |  |  |  | | |  | | |  | | |  | | |  | | |  | | |  | | |  | | |  | | |
|  |  |  |  |  | Workers | | | | Females | | | 0.26 | | |  | | |  |  |  |  |  |  | | |  | | |  | | |  | | |  | | |  | | |  | | |  | | |  | | |
|  | PB | No  p = 0.003 | | | Males | | | | Workers | | | 0.08 | | |  | | | Males < 0.01 Workers = 0.01 Females = 0.04 | |  | | |  | | |  | | |  | | |  | | |  | | |  | | |  | | |  | | |  | | |
|  |  |  |  |  | Females | | | | Males | | | 0.007 | | |  | | |  |  |  |  |  |  | | |  | | |  | | |  | | |  | | |  | | |  | | |  | | |  | | |
|  |  |  |  |  | Workers | | | | Females | | | 0.0006 | | |  | | |  |  |  |  |  |  | | |  | | |  | | |  | | |  | | |  | | |  | | |  | | |  | | |
|  | NO | Yes  p = 0.06 | | |  | | | |  | | |  | | | Yes p = 0.04 | | | Males = 0.01 Workers = 0.01 Females =0.36 | | Yes p = 7.76 x 10^-10^ | | | Males | | | Workers | | | 2.93 x 10^-10^ | | | -4.39 | | | -4.81 | | | -4.71 | | | 2.67 | | | 0.48 | | | 0.79 | | |
|  |  |  |  |  |  | | | |  | | |  | | |  |  |  |  |  |  |  |  | Females | | | Males | | | 1.42 x 10^-4^ | | |  |  |  |  |  |  |  |  |  |  |  |  |  |  |  |  |  |  |
|  |  |  |  |  |  | | | |  | | |  | | |  |  |  |  |  |  |  |  | Workers | | | Females | | | 0.06 | | |  |  |  |  |  |  |  |  |  |  |  |  |  |  |  |  |  |  |
|  | CX | Yes  p = 0.2 | | |  | | | |  | | |  | | | Yes p = 0.03 | | | Males < 0.01 Workers < 0.01 Females = 0.12 | | Yes p = 6.01 x 10^-14^ | | | Males | | | Workers | | | 6.77 x 10^-15^ | | | -3.23 | | | -3.55 | | | -3.47 | | | 2.1 | | | 0.57 | | | 0.84 | | |
|  |  |  |  |  |  | | | |  | | |  | | |  |  |  |  |  |  |  |  | Females | | | Males | | | 1.22 x 10^-6^ | | |  |  |  |  |  |  |  |  |  |  |  |  |  |  |  |  |  |  |
|  |  |  |  |  |  | | | |  | | |  | | |  |  |  |  |  |  |  |  | Workers | | | Females | | | 0.11 | | |  |  |  |  |  |  |  |  |  |  |  |  |  |  |  |  |  |  |
